# Quantitative phosphoproteomics uncovers dysregulated kinase networks in Alzheimer’s disease

**DOI:** 10.1101/2020.08.18.255778

**Authors:** Nader Morshed, Meelim Lee, Felicia H. Rodriguez, Douglas A. Lauffenburger, Diego Mastroeni, Forest White

**Affiliations:** Department of Bioengineering, Massachusetts Institute of Technology, Cambridge, MA, 02139, USA; Koch Institute for Integrative Cancer Research, Massachusetts Institute of Technology, Cambridge, MA 02139, USA; Department of Chemical and Materials Engineering, New Mexico State University, Las Cruces, NM 88003, USA; Arizona State University-Banner Neurodegenerative Disease Research Center, Tempe, Arizona 85287, USA; Center for Precision Cancer Medicine, Massachusetts Institute of Technology, Cambridge, MA 02139, USA

## Abstract

Alzheimer’s disease (AD) is a form of dementia characterized by amyloid-β plaques and Tau neurofibrillary tangles that progressively disrupt neural circuits in the brain. The signaling networks underlying the pathological changes in AD are poorly characterized at the level phosphoproteome. Using mass spectrometry, we performed a combined analysis of the tyrosine, serine, and threonine phosphoproteome, and proteome of temporal cortex tissue from AD patients and aged matched controls. We identified several co-correlated peptide modules that were associated with varying levels of phospho-Tau, oligodendrocyte, astrocyte, microglia, and neuronal pathologies in AD patients. We observed phosphorylation sites on kinases targeting Tau as well as other novel signaling factors that were correlated with these peptide modules. Finally, we used a data-driven statistical modeling approach to identify individual peptides and co-correlated signaling networks that were predictive of AD histopathologies. Together, these results build a map of pathology-associated phosphorylation signaling events occurring in AD.

## Introduction

Alzheimer’s disease (AD) is a neurodegenerative disease commonly associated with aging. AD has an immense disease burden, with an estimated 50 million cases worldwide and a total economic cost of more than $500 billion per year^1^. Despite numerous clinical trials, a limited understanding of disease pathogenesis has hindered the development of effective therapies. Amyloid beta (Aβ) plaques have been identified as a hallmark pathology of AD that develops early on, followed by accumulation of hyper-phosphorylated microtubule-associated protein Tau (MAPT) and eventual neurodegeneration^2^. Throughout AD, cellular signaling through protein phosphorylation events mediates Aβ and MAPT production and cytotoxic mechanisms as well as other stress-responses^3^. Previous work has identified GSK3β, CDK5, Protein Kinase A, CaMKII, Fyn, and members of the BRSK, MARK, MAPK, and Casein Kinase families as enzymes that can phosphorylate MAPT and promote its aggregation^4–7^. Other dysregulated kinases include PKCα as a mediator of synaptic defects^8,9^, JAK as an activator of inflammation^10,11^, and mTOR as a regulator of neuronal homeostatic adaption^12^. Targeting some of these kinases has shown positive effects in AD models^8,13,14^, suggesting that modulating cellular signaling through protein phosphorylation may be a viable treatment strategy for AD. However, there have been few studies mapping these signaling events to stages of AD progression using knowledge of the phosphoproteome.

Tools to measure the proteome and phosphoproteome have been developed using mass spectrometry (MS)^15–20^. Previous studies have analyzed the phosphoproteome of AD mouse models^21–23^ as well as the global phosphoserine/phosphothreonine (pSer/pThr) phosphoproteome and global proteome of post-mortem AD brain tissue^21,24–30^. However, these studies were limited by an incomplete characterization of the phosphoproteome that did not include critical phosphotyrosine (pTyr) sites that mediate the activity of many known kinases. Many of these studies were also limited to a small number of patients or pooled samples together, which hinders the development of a quantitative model of disease progression. We aimed in this study to integrate pTyr, deep pSer/pThr phosphoproteome, and deep proteome measurements of AD and age-matched brain tissues over a range of disease stages. Our data collection strategy allowed us to associate phosphoproteome changes with direct measurements of MAPT phosphorylation, cell-type specific markers, and clinical histopathology measurements.

Co-correlation analysis of this dataset identified several clusters that were enriched for neuronal and glial-specific proteins; these clusters were associated with Tau pathology, oligodendrocyte changes,and neurodegeneration. Our analysis recaptured the links between Tau pathology and several known MAPT kinases and highlighted kinases and signaling factors that were previously unreported to be associated with AD. These factors include creatine kinase B-type (CKB), cyclin-dependent kinases (CDKs)CDKL5, CDK16, CDK17, and the tyrosine kinases SRC, DDR1, EGFR, and PTK2. Integrative analysis of phosphoproteomic and proteomic data with patient histopathology measurements highlight intriguing association between many of these signaling factors and AD pathologies, providing new insight into disease progression. Together these data underscore the value of phosphoproteomics technologies in building a quantitative understanding of disease etiology at a systems-scale.

## Methods

### Human brain samples

Samples of human middle temporal gyrus (MTG) were secured from AD or neurologically normal, non-demented (ND) elderly control brains obtained from rapid autopsy at the Banner Sun Health Research Institute Tissue Bank (BSHRI)^31,32^. Cognitive status of all cases was evaluated antemortem by board-certified neurologists, and postmortem examination by a board-certified neuropathologist resulting in a consensus diagnosis using standard NIH AD Center criteria for AD or ND. The AD and ND groups were well matched for age, sex and PMI. ND control cases: ND Braak I n=5, ND Braak IV n=18 (23 total); Age: 84.1 ± 7.1 years; Gender: 12 females and 11 males; Postmortem interval (PMI: 2 hours 58 minutes +/− 75 min. AD cases: AD Braak IV n=10, AD Braak V n=8, AD Braak VI n=7; Age: 84.3 +/− 7.9 years; Gender: 11 females and 14 males; PMI: 3 hours 23 min +/− 180 min. ND patients were classified as APP^+^ if they had a Plaque density value that was not “zero”. The full list of each ID, sex, age, and diagnostic histopathology measurements for patients is provided in **Supplementary Table 1**.

### Sample Processing for Proteomics

MS-based analysis of pTyr, pSer/pThr, and protein expression were conducted using a standard analysis pipeline developed in the White lab and used in numerous studies and publications^19,33–36^. 20 - 60 mg of frozen tissues for proteomics analysis were transferred into 3 mL ice-cold 8M Urea and immediately homogenized to denature proteins and preserve physiological signaling. Lysates were then centrifuged at 4000xg for 30 minutes to clear lysates of lipids and then assayed by BCA to determine protein concentration. Lysates were then treated with 10 mM DTT for 1 hour at 56°C, followed by 55 mM iodoacetamide for 1 hour at room temperature, rotating in the dark. Samples were diluted by adding 8 mL Ammonium Acetate pH 8.9 and then digested at room temperature overnight on a rotator with trypsin at a 1:50 ratio of trypsin to total protein mass. This reaction was quenched with 1 mL 99.99% Acetic Acid and samples were desalted using Sep-Pak Plus cartridges. Peptides were then dried to half volume in a speed-vac and post-cleanup concentrations were determined using a colorimetric peptide assay. Peptides were then divided into 130 μg aliquots that were snap-frozen and lyophilized. Peptides were then labeled with 10-plex isobaric Tandem Mass Tags (TMT) using a reduced volume protocol^37^. Labeled peptide samples were then dried down in a speed-vac overnight.

### Peptide Fractionation and Phosphopeptide Enrichment

Phospho-enrichment experimental design is pictured in **Figure 1a**. Tyrosine phosphorylated peptides were enriched by a two-step process consisting of an immunoprecipitation (IP) with multiple pan-specific anti-phosphotyrosine antibodies (4G10, PT66) followed by immobilized metal affinity chromatography (IMAC) using hand-packed micro-columns as we have previously described^33,38–41^ as well as with commercial IMAC spin-columns. IP supernatants were subjected to a second round of IP enrichment with anti-MAPK-CDK phospho substrate antibody followed by IMAC cleanup using commercial spin-columns. Small amounts of each IP supernatant were diluted in 0.1% acetic acid for global protein expression profiling. IP supernatants were then divided across 80 fractions using high pH reverse phase chromatography on a ZORBAX C18 column. Fractions were concatenated into 20 tubes and dried down. Each fraction was then enriched using commercial Fe-NTA columns. Small amounts of each fraction were diluted in 0.1% acetic acid for deep proteome profiling.

**Figure 1.**
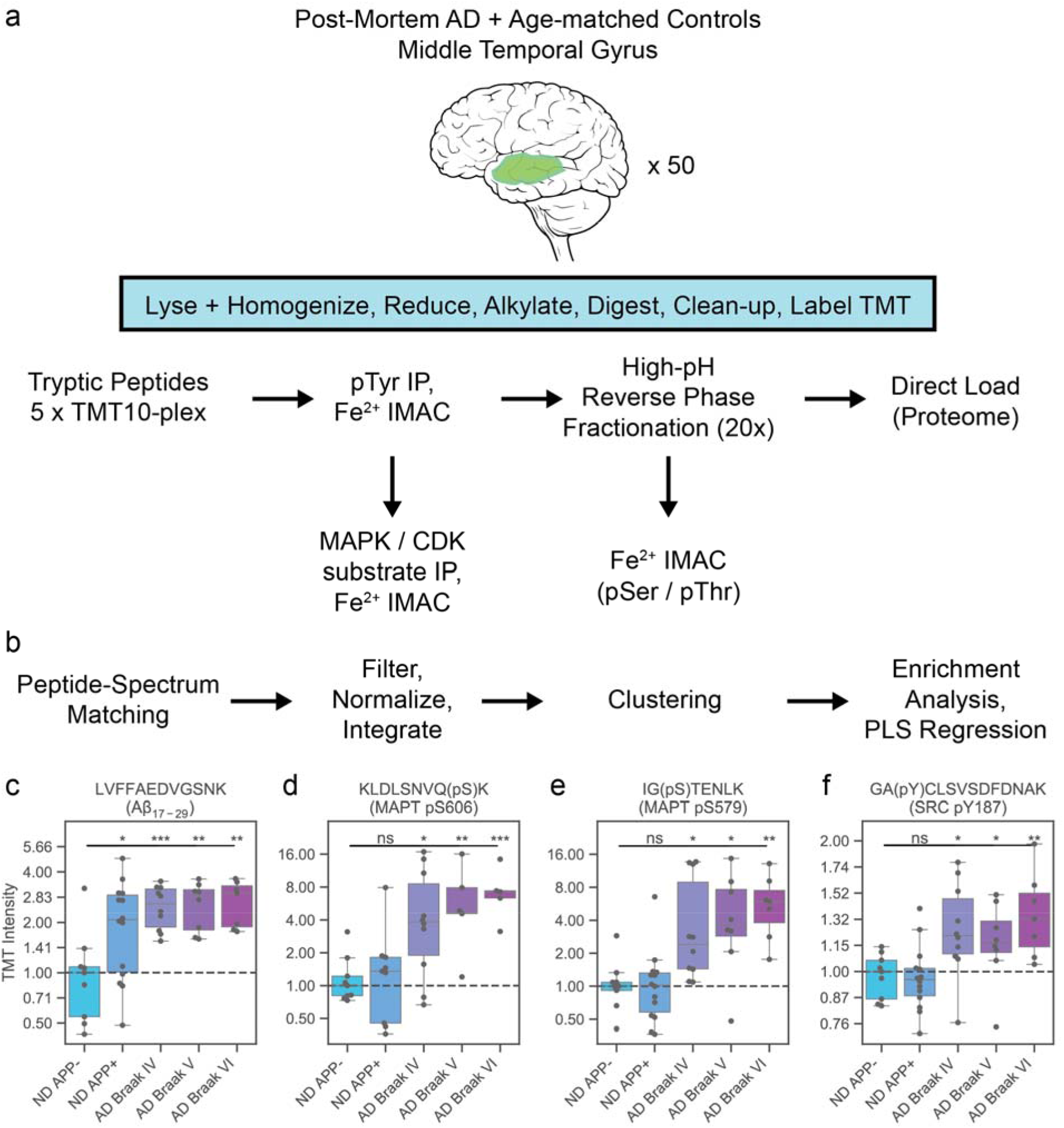
Combined phosphoproteomics and proteomics analysis captures molecular signaling profile of Alzheimer’s disease. (a) Workflow describing proteomics sample processing and peptide enrichment. Frozen patient tissue was lysed and homogenized to extract proteins and then reduced, alkylated, and digested into short tryptic peptides. Proteolytic digests were then labeled with TMT10-plex for multiplexed quantification. pTyr peptides were pulled down by antibody and cleaned up using IMAC. IP supernatants were then IP’d for MAPK-CDK substrates and then fractionated offline at high-pH into 20 concatenated fractions. Fractions were then direct-loaded onto an autosampler for global proteomics and then enriched for global pSer/pThr peptides using IMAC. All enriched peptides were then analyzed on a Q Exactive mass spectrometer to identify and quantify the relative abundance across samples. (b) Raw mass spectrometry data was searched using ProteomeDiscoverer and MASCOT to generate peptide-spectrum matches (PSMs). PSMs were filtered to remove low-quality matches and TMT abundances were normalized to compare across 10-plex runs. PSMs matching the same peptide sequence were then merged together to generate a final peptide abundance matrix. Peptide data were analyzed via K-means clustering and combined with clinical pathology assessments using PLSR modeling to identify covarying data trends. Final peptide modules were used for gene-set, kinase-substrate set, phosphorylation-motif, cell-type and PPI-enrichment analyses to identify network-level trends. (c-f) Selected trends are shown for peptides matched to (c) Aβ_17-28_, (d) MAPT pS606, (e) MAPT pS579, and (f) SRC pTyr187. Each plot shows peptide fold changes for integrated TMT values. Values are normalized to the median of ND APP-patients. Stars indicate t-test values for each group with ND APP-patients. ns, not significant (p > 0.05); **p < 1e-2; ***p < 1e-3; ****p < 1e-4.

For commercial spin-columns, Thermo High-Select Fe-NTA columns were washed twice with 200 μL Binding/Wash Buffer and beads were resuspended in 25 μL Binding/Wash Buffer. Peptides were eluted from IP beads using two rounds of 10 min washes with 25 μL 0.2% trifluoroacetic acid (TFA) using the same pipette tip to transfer eluates directly onto Fe-NTA beads. Phosphopeptides were incubated with beads for 30 minutes with gentle tapping. Beads were then washed twice with 200 μL Binding/Wash Buffer, and once with 200 μL LC-MS water. Phosphopeptides were eluted into BSA-coated microcentrifuge tubes using two 20 μL washes of Phosphopeptide Elution Buffer. Samples were brought down to 1-5 μL volume by speed-vac for ~20 minutes and then 10 μL 0.1% acetic acid with 2% acetonitrile was added. Samples were loaded directly onto a hand-packed, BSA-conditioned 10 cm analytical column with 5 μm C18 beads. Columns were rinsed with 0.1% acetic acid to remove excess salts and then analyzed by liquid chromatography-tandem mass spec (LC-MS/MS) on QExactive HF-X Orbitrap mass spectrometer.

For LC analysis, Agilent 1100 Series HPLCs were operated at 0.2 mL/min flow rates with a pre-column split to attain nanoliter flow rates through the analytical column and nano-ESI tip. Peptides were eluted with increasing concentrations of buffer B (70% acetonitrile, 0.1% acetic acid) using the gradient settings: 0-13% (10 min), 13-42% (95 min), 42-60% (10 min), 60-100% (7 min), 100% hold (6 min), 100-0% (2 min). Global phosphoproteome and proteome fractions were analyzed using an EASY-nLC nano-flow UHPLC. Fractions were eluted using the gradient settings: 0-10% (10 min), 10-30% (100 min), 30-40% (14 min), 40-60% (5 min), 60-100% (2 min), hold 100% (10 min), 100-0% 2 min. Peptides were ionized by electrospray ionization (ESI) at 1 - 3 kV.

Peptides were analyzed by LC-MS/MS on a QExactive Plus and QExactive HF-X Orbitrap mass spectrometer operating in data-dependent mode acquiring MS scans and HCD MS/MS fragmentation spectra. Ions with charge >1 were dynamically selected for fragmentation using a top-20 untargeted method with an exclusion time of 30s and ion isolation window of 0.4 Da. The maximum injection time and ACG targets were set to 50 ms and 3e6 respectively. MS scans were captured with resolution = 60,000 and MS2 scans with resolution = 45,000. Peptides were fragmented with the HCD normalized collision energy set to 33%. Protein expression profiling was performed on LTQ Orbitrap or QExactive Plus instruments.

### Data Processing and Analysis

Peptide identification and quantification was performed using Proteome Discoverer and MASCOT. Raw files were searched against the ‘SwissProt_2020_02.fasta’ for tryptic peptides from *Homo sapiens* with ≤2 missed cleavages. Precursor and fragment ions were matched with a 10 ppm and 20 mmu mass tolerances respectively. Variable modifications were included for phosphorylation (Ser, Thr, Tyr), oxidation (Met), and TMT 10plex (Lys, N-term) and fix modifications were set for carbamidomethyl (Cys). False discovery rates for peptide-spectrum matches (PSMs) were estimated using the Percolator module and post-translational modification (PTM) localizations were calculated using ptmRS^42^. ptmRS operated with a peak depth of 15. For pTyr analyses, a diagnostic ion of 216.04 with a peak depth of 15 was used. For MAPK/CDK substrate, pSer/pThr, and proteome analyses, a neutral loss of 98 (-H_3_PO_4_) was included for pSer/pThr ions.

Searched .msf files were then imported into Python and processed using pyproteome (https://github.com/white-lab/pyproteome). PTMs were localized using a ptmRS confidence cutoff of 0.75. PSMs were filtered for ion score ≥15, isolation interference ≤30, median TMT signal ≥1500, and Percolator FDR ≤1e-2. TMT quantification data was then normalized using a iterative fitting procedure based on the CONSTrained STANdardization algorithm^43^. In this procedure, the matrix of filtered TMT quantification values was iteratively adjusted such that rows were mean-centered around 1 and columns were mode-centered around 1. Mode centers were calculated by fitting a gaussian KDE distribution (scipy.stats.kde.gaussian_kde) to column intensities and finding the max value of each probability density function. Duplicate PSMs were then grouped for final quantification using a weighted mean function: 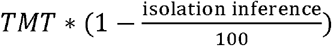 for the TMT intensities and isolation interference value quantified by Proteome Discoverer for each PSM. Final relative TMT intensities were normalized to ND APP^−^ control samples across TMT 10-plex analyses. Data clustering, statistical analysis, and figure generation was performed using *sci-kit learn*^44^, *scipy*^45^, *numpy*^46^, *pandas*^47^, *seaborn*^48^, and *matplotlib*^49^.

### Clustering and regression analysis

Clustering analysis was performed on the unique peptide abundance matrix using rows with no missing values for peptides that matched ≤4 proteins. Mini-Batch K-means clustering was performed using scikit-learn’s ‘MiniBatchKMeans’ function. A cluster number of 100 was selected based on an analysis of cluster membership properties. Cluster centroids were calculated from the mean log_2_ value of all peptides assigned to a cluster.

For partial least squares regression (PLSR) analysis, the X matrix was generated from all cluster centroid log_2_ values and the Y matrix was generated from histopathology measurements. Select histopathology measurements were translated into discrete numbers that were then z-scored. 2-component PLSR models were generated using pyls, a Python implementation of the SIMPLS algorithm. A K-fold cross validation procedure was run for each histopathology variable with splits=5 and repeats=100. R^2^ values were equal to the Spearman ρ^2^ between the true and predicted Y vectors for the training datasets. Q^2^ values were similarly calculated from the true and predicted Y vectors from the test datasets. A final 2-component PLSR model was calculated from non-redundant histopathology variables with Q^2^ >.3. PLSR model variables are related by the equation X * X weights = X scores and Y_approximate_= Y scores_truncated_ * Y loadings_truncated_, where truncated indicates that the model is being truncated at 2 latent variables.

Explained variance for Y matrices was calculated as the ratio of the sum of squares of the Y_loadings_ matrix to the sum of squares of the Y matrix. A similar formula was used for the ratio of the X_loadings_ matrix (calculated from the pseudoinverse of the X_weights_) to the X matrix.

To select strongly-associated variables, Variable Importance in Projection (VIP) scores were calculated using the previously-described formulae^50^:

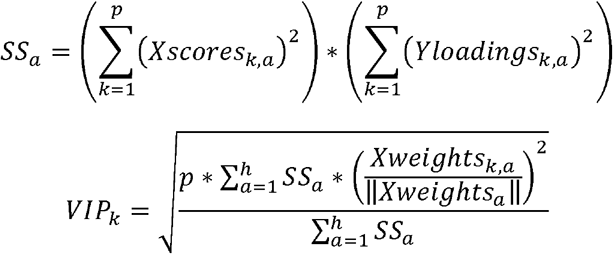

Where a is the a^th^ component, k is the k^th^ predictor variable, p is the number of predictor variables, h is the number of components.

Correlation cutoffs were estimated by generating a null distribution from all peptides quantified in ≥15 samples. For each cluster, we calculated Spearman’s correlation between peptides and scrambled cluster centroid values. We repeated this procedure 100 times and the 99^th^ percentile value was estimated from all calculated Spearman ρ and Spearman p-values. The final ρ and p-value cutoffs of 0.47 and 5e-3 were selected as cutoffs that matched less than 0.24% of all peptides for any of our six named cluster centroids. Spearman partial correlation coefficients were calculated using ‘pingouin.pairwise_corr’. Receiver operating characteristic (ROC) curves were generated using ‘sklearn.metrics.roc_curve’.

### Cell type enrichment and composition analysis

Gene cell-type predictions were estimated using a cell-type specific sequencing atlas^51,52^. Cell expression values were extracted from Table S4 using the following columns: ‘Astrocyte’: [‘8yo’, ‘13yo’, ‘16yo’, ‘21yo.1’, ‘22yo.1’, ‘35yo’, ‘47yo’, ‘51yo’, ‘53yo’, ‘60yo’, ‘63yo - 1’, ‘63yo - 2’], ‘Neuron’: [’25 yo’], ‘OPC’: [‘22yoGC’, ‘63yoGC - 1’, ‘63yo GC - 2’, ‘47yoO4’, ‘63yoO4’], ‘New Oligodendrocytes’: [‘22yoGC’, ‘63yoGC - 1’, ‘63yo GC - 2’, ‘47yoO4’, ‘63yoO4’], ‘Myelinating Oligodendrocytes’: [‘22yoGC’, ‘63yoGC - 1’, ‘63yo GC - 2’, ‘47yoO4’, ‘63yoO4’], ‘Microglia’: [‘45yo’, ‘51yo.1’, ‘63yo’], ‘Endothelia’: [‘13yo.1’, ‘47yo.1’]. Genes were calculated as being enriched in a given cell type if they met the criteria 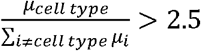, where μ is the mean FPKM value from collections of columns for each cell type. OPC and New Oligodendrocytes were excluded from the enrichment calculation to avoid exclusion of pan-oligodendrocyte genes, only ‘Myelinating Oligodendrocytes’ was used for the displayed ‘Oligodendrocyte’ category.

For cell type composition analysis, peptides were processed into a list of unique cell type-enriched genes with at least one PSM (modified or unmodified) that was strongly correlated with each centroid (Spearman ρ > .7; Spearman p-value < 1e-3; # patients ≥15). Phosphorylation composition was performed similarly by mapping each unique peptide to ‘pY’ if it contained at ≥1 pTyr modification, ‘pST’ if it contained ≥1 pST modification, or ‘no phospho’ if it was not phosphorylated. Cluster cell type and phosphorylation significance was calculated using scipy’s ‘chi2_contingency’ comparison between each cluster centroid and the background dataset of peptides (# patients ≥15).

### Kinome analysis

Kinase phosphopeptides that were quantified in ≥15 patients and were significantly correlated with Tau or Oligo were used for kinome analysis. We generated vectors for each peptide that were equal to the Spearman partial ρ between that peptide and each cluster centroid (ρ_Tau_, ρ_Oligo_), with the alternative cluster centroid as the covariate. Correlation vectors were filtered to have angles between −25° and 110°, corresponding to the range of kinase phosphosites that were significantly correlated with Tau or Oligo clusters. Phosphopeptides mapping to more than one kinase are used to color all nodes. For each kinase, the phosphopeptide with the largest vector magnitude was used to calculate node color and size. Kinome maps were generated using Coral^53^ with nodes and branches colored by the correlation vector angle.

### Binomial enrichment analysis

Binomial enrichment analysis of pTyr, pSer/pThr, and protein data subsets was performed using gene sets derived from published proteomics data^26^ and single-nuclei RNA-seq data^54^. Peptides were processed into a background collection of unique genes that were enriched in any gene set. Only peptides that mapped to a single protein were used. Foreground collections were generated from peptides that had spearman correlation >.7 with cluster centroids. Log odds enrichment (LOE) values were calculated as – 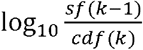, with k = the count of genes in a given gene set from the foreground collection. Survival functions (sf) and cumulative distribution functions (cdf) were calculated using scipy’s hypergeometric distribution model (scipy.stats.hypergeom), with: n = the total number of foreground genes; K = the count of genes in the gene set; and N=the total number of background genes.

### Phosphosite Enrichment Analysis

Phosphosite enrichment analysis (PSEA) were performed using the procedure outlined in a previous publication^55^. We used a custom phosphosite database derived from Phosphosite Plus (PSP)^56^. Kinase-substrate mappings were downloaded from PSP using information for all species (Kinase_Substrate_Dataset.gz), and then re-mapped to human phosphosites using homology information from PSP (Phosphorylation_site_dataset.gz). BRSK1 and BRSK2 phosphosite sets were generated from a recent phosphoproteomics analysis^57^ using the table of filtered and imputed abundance values with a MaxQuant localization probability > 0.6, log_2_ Fold Change of 1.5, and 2-sample T-test p-value < 1e-2 for log_2_ abundance values.

Phosphosite changes were estimated from the median value of redundant phosphopeptides. Spearman correlations with cluster centroids were z-scored and phosphosites were rank ordered. Enrichment scores were calculated using the integral of a running-sum statistic (exponential weight = 0.75) that was increased for each gene or site contained within a given set and decreased for each gene not contained in that set. Enrichment scores for 1000 matrices with scrambled rows were generated and used to calculate empirical p-values, q-values, and normalized enrichment scores (NES). Only phosphosite sets with a minimum overlapping set size of 20 were scored.

### Mouse Phosphoproteome Analysis

The phosphoproteome of AD mouse models was downloaded from a recent paper that was collected using a similar TMT quantification and phosphopeptide enrichment strategy (ProteomeXchange identifier: PXD018757)^22^. Mouse phosphosites were categorized as upregulated (Fold Change > 1.25; p < 1e-2) or downregulated (Fold Change < .8; p < 1e-2) for all tissues and disease timepoints in each model (CK-p25: 2wk Hippocampus + Cortex, 7wk Hip; 5XFAD: 9-10mo Hippocampus + Cortex; Tau P301S: 4+6 mo Hippocampus). Phosphosites were mapped from mouse to human using the most recent version of ‘Phosphorylation_site_dataset.gz’ from PSP^56^. Missing phosphosite mappings were automatically generated by matching a 9-mer around the mouse phosphosite to corresponding human protein sequence using Python’s ‘difflib.SequenceMatcher’ function. Phosphosite overlap was calculated between each mouse model and the human phosphosites that were correlated with each cluster centroid (Spearman ρ > .47; Spearman p-value < 5e-3; # patients ≥15). Phosphoproteins were counted if they had at least one homologous phosphopeptide that was associated with a cluster.

### Data and Code Availability

The mass spectrometry proteomics data have been deposited to the ProteomeXchange Consortium via the PRIDE^58^ partner repository with the dataset identifier PXD020087 and 10.6019/PXD020087.

The proteomics data integration software is available on GitHub (https://github.com/white-lab/pyproteome). This repository also includes detailed software for concatenating peptide fractions on a Gibson FC 204 Fraction Collection (https://github.com/white-lab/fc-cycle), and an updated tool for validating PSMs (https://github.com/white-lab/CAMV). Sources for other code used in this study are indicated in the Resources Table.

### Resources Table

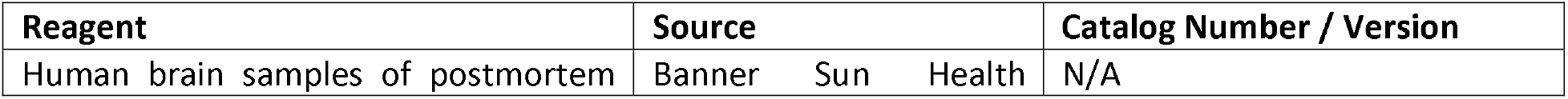
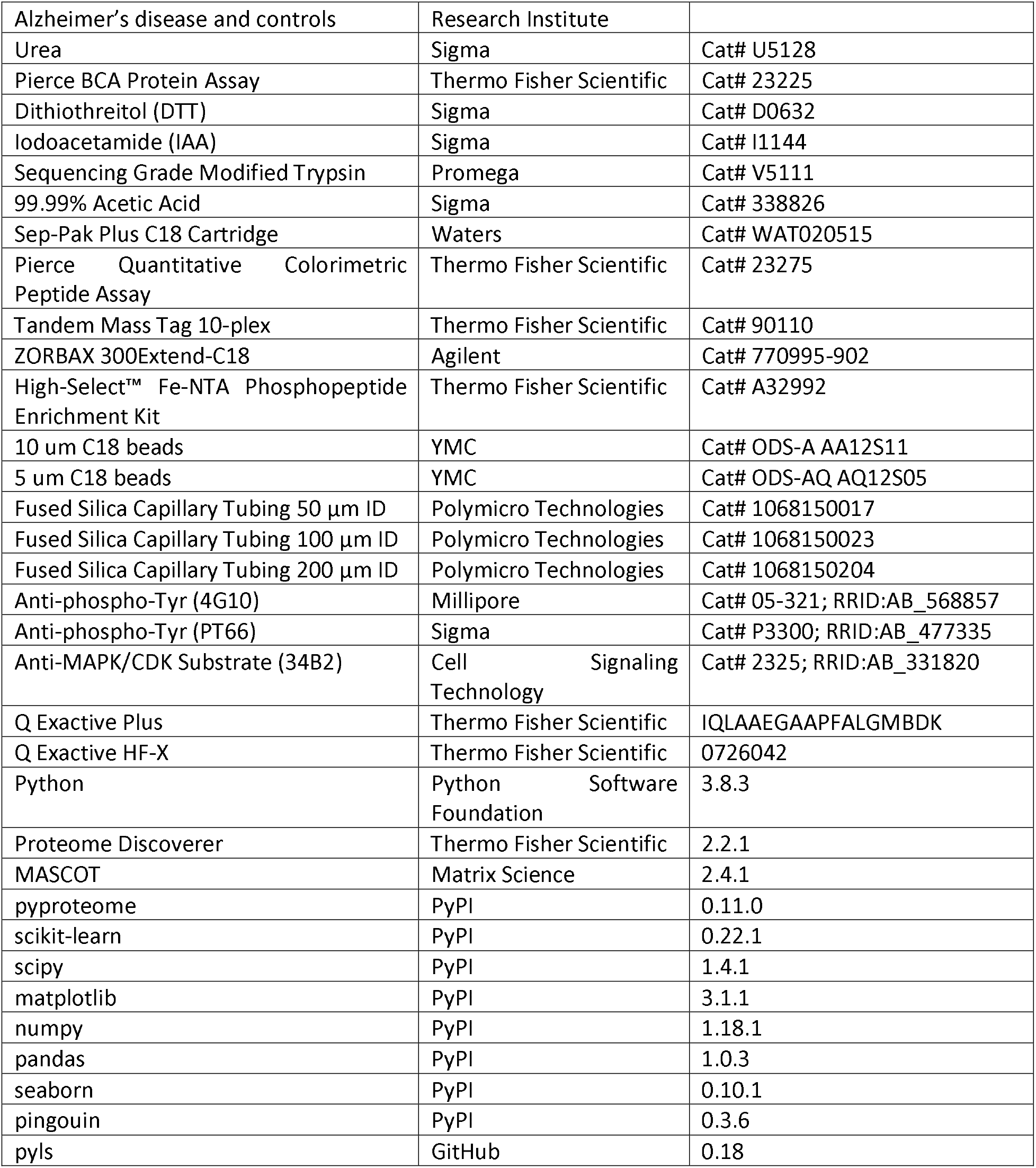

| Reagent | Source | Catalog Number / Version |
| --- | --- | --- |
| Human brain samples of postmortem | Banner Sun Health | N/A |
| Alzheimer's disease and controls | Research Institute |  |
| Urea | Sigma | Cat# U5128 |
| Pierce BCA Protein Assay | Thermo Fisher Scientific | Cat# 23225 |
| Dithiothreitol (DTT) | Sigma | Cat# D0632 |
| Iodoacetamide (IAA) | Sigma | Cat# I1144 |
| Sequencing Grade Modified Trypsin | Promega | Cat# V5111 |
| 99.99% Acetic Acid | Sigma | Cat# 338826 |
| Sep-Pak Plus C18 Cartridge | Waters | Cat# WAT020515 |
| Pierce Quantitative Colorimetric Peptide Assay | Thermo Fisher Scientific | Cat# 23275 |
| Tandem Mass Tag 10-plex | Thermo Fisher Scientific | Cat# 90110 |
| ZORBAX 300Extend-C18 | Agilent | Cat# 770995-902 |
| High-Select™ Fe-NTA Phosphopeptide Enrichment Kit | Thermo Fisher Scientific | Cat# A32992 |
| 10 µm C18 beads | YMC | Cat# ODS-A AA12S11 |
| 5 µm C18 beads | YMC | Cat# ODS-AQ AQ12S05 |
| Fused Silica Capillary Tubing 50 µm ID | Polymicro Technologies | Cat# 1068150017 |
| Fused Silica Capillary Tubing 100 µm ID | Polymicro Technologies | Cat# 1068150023 |
| Fused Silica Capillary Tubing 200 µm ID | Polymicro Technologies | Cat# 1068150204 |
| Anti-phospho-Tyr (4G10) | Millipore | Cat# 05-321; RRID:AB_568857 |
| Anti-phospho-Tyr (PT66) | Sigma | Cat# P3300; RRID:AB_477335 |
| Anti-MAPK/CDK Substrate (34B2) | Cell Signaling Technology | Cat# 2325; RRID:AB_331820 |
| Q Exactive Plus | Thermo Fisher Scientific | IQLAAEGAAPFALGMBDK |
| Q Exactive HF-X | Thermo Fisher Scientific | 0726042 |
| Python | Python Software Foundation | 3.8.3 |
| Proteome Discoverer | Thermo Fisher Scientific | 2.2.1 |
| MASCOT | Matrix Science | 2.4.1 |
| pyproteome | PyPI | 0.11.0 |
| scikit-learn | PyPI | 0.22.1 |
| scipy | PyPI | 1.4.1 |
| matplotlib | PyPI | 3.1.1 |
| numpy | PyPI | 1.18.1 |
| pandas | PyPI | 1.0.3 |
| seaborn | PyPI | 0.10.1 |
| pingouin | PyPI | 0.3.6 |
| pyls | GitHub | 0.18 |

## Results

### Proteomic and phosphoproteomic analyses identify molecular hallmarks of AD

In order to analyze the AD proteome and phosphoproteome, we generated proteolytic peptide digests of medial temporal gyrus (MTG) cortical brain tissue from patients with late-onset AD and age-matched non-diseased (ND) controls. Previous work has shown that the proteome of this region is affected by AD in a similar manner to other brain regions^26^. We used isobaric tandem mass tags (TMT) to label peptides, allowing us to perform multiplexed relative quantification of peptide abundances across samples. Using a sample processing and peptide enrichment scheme outlined in **Figure 1a**, we quantified relative changes in the proteome and phosphoproteome of 48 patient samples (AD n=25; ND n=23; Age=70-90; Female=23; Male=25; **Supplementary Data 1**). We performed an analysis of the low-abundance pTyr phosphoproteome as well as a deep fractionated analysis of pSer/pThr phosphoproteome and proteome to generate a large dataset of peptide-spectrum matches (PSMs). By integrating TMT abundances across all runs, we constructed a matrix of relative peptide abundances in each patient that we could use to interrogate signaling network changes. This strategy led to the identification and quantification of 88,419 unique peptides (1.5% pTyr, 16% pSer/pThr, **Supplementary Data 2**), of which 58% of all peptides were identified in at least 15 patient samples (≥2 10-plex analyses) **(Supplementary Figure 1a-d)**. To assess the reproducibility of our quantification method, we compared peptide abundances between technical replicates of TMT10-plex analyses. To assess biological variability, we additionally analyzed two independent brain samples in separate TMT10-plex analyses for two patients. We found that peptide abundances were well correlated between samples for both technical and biological replicates **(Supplementary Figure 1e-h)**. With a sensitive and quantitative data analysis strategy, we next examined the molecular features within individual patient proteomes and phosphoproteomes.

Our proteomics dataset successfully recapitulates hallmark AD pathologies, including a tryptic peptide mapping to Aβ_17-28_ (LVFFAEDVGSNK) that was upregulated in some ND patients and plateaued in AD Braak IV-VI cases **(Figure 1c)**. Additionally, we identified phosphopeptides from MAPT (KLDLSNVQsK: pS606; IGsTENLK: pS579) that were increased in AD patients **(Figure 1d, 1e)**. Within the pTyr dataset, we observed increased phosphorylation of the non-receptor tyrosine kinase SRC (GAycLSVSDFDNAK: pY187; **Figure 1f**). We note that these peptides are upregulated at different Braak stages and have different ranges of variability, reflecting the complex etiology of AD combined with other potential patient comorbidities. Together, this dataset allows us to infer aspects of disease pathology and link those measurements to the phosphosites regulating kinase activity in a given tissue sample.

### Clustering analysis identifies Tau and cell-type peptide modules

In order to identify major trends of covariation in our dataset that may be reflective of different types of pathology in each patient, we applied clustering analysis to extract co-correlated peptide changes. We performed K-means clustering on the 8,706 peptides that were quantified in all 48 patients (1.6% pTyr, 13% pSer/pThr; **Supplementary Data 3**). We selected a cutoff of n=100 clusters to balance the intra-cluster correlation value with the number of redundant clusters **(Supplementary Figure 2a-2c)**. Of the 100 clusters, we manually assigned identities to six clusters that explained 73% of all cluster variance. “Tau” is a cluster that includes several MAPT peptides that were upregulated in AD patients; rank-ordering the patient samples based on the signal intensity of the Tau cluster effectively separates the ND and AD patients, albeit with a few exceptions **(Figure 2a)**. “Oligo”, “Astro”, and “Micro” are three clusters containing proteins primarily expressed in oligodendrocytes (MBP, PLP1), astrocytes (GFAP), and microglia (GNA13) respectively **(Figure 2b-d)**. “Neuro” and “pNeuro” are two clusters containing proteins (SYT1, SYN1, SNAP25) and phosphopeptides (CAMKK1, GSK3A, CASKIN1) involved in neuronal synaptic signaling; both of these clusters were downregulated in subsets of AD patients **(Figure 2e, 2f)**. We examined the composition of predicted cell types for peptides that were strongly correlated with each cluster’s centroid and found that the Oligo, Astro, Micro, and Neuro clusters were significantly enriched for each respective cell type **(Figure 2g)**. In addition, the Tau and pNeuro clusters contained an increased fraction of pSer/pThr peptides compared to the full dataset **(Supplementary Figure 2d)**. The cluster centroids for Tau and Oligo were independent (correlation ≈ 0), while Astro and Micro were correlated with one another as well as with Tau and Oligo **(Figure 2h)**. In contrast, the Neuro cluster was most strongly anti-correlated with Oligo and Micro **(Figure 2h)**.

**Figure 2.**
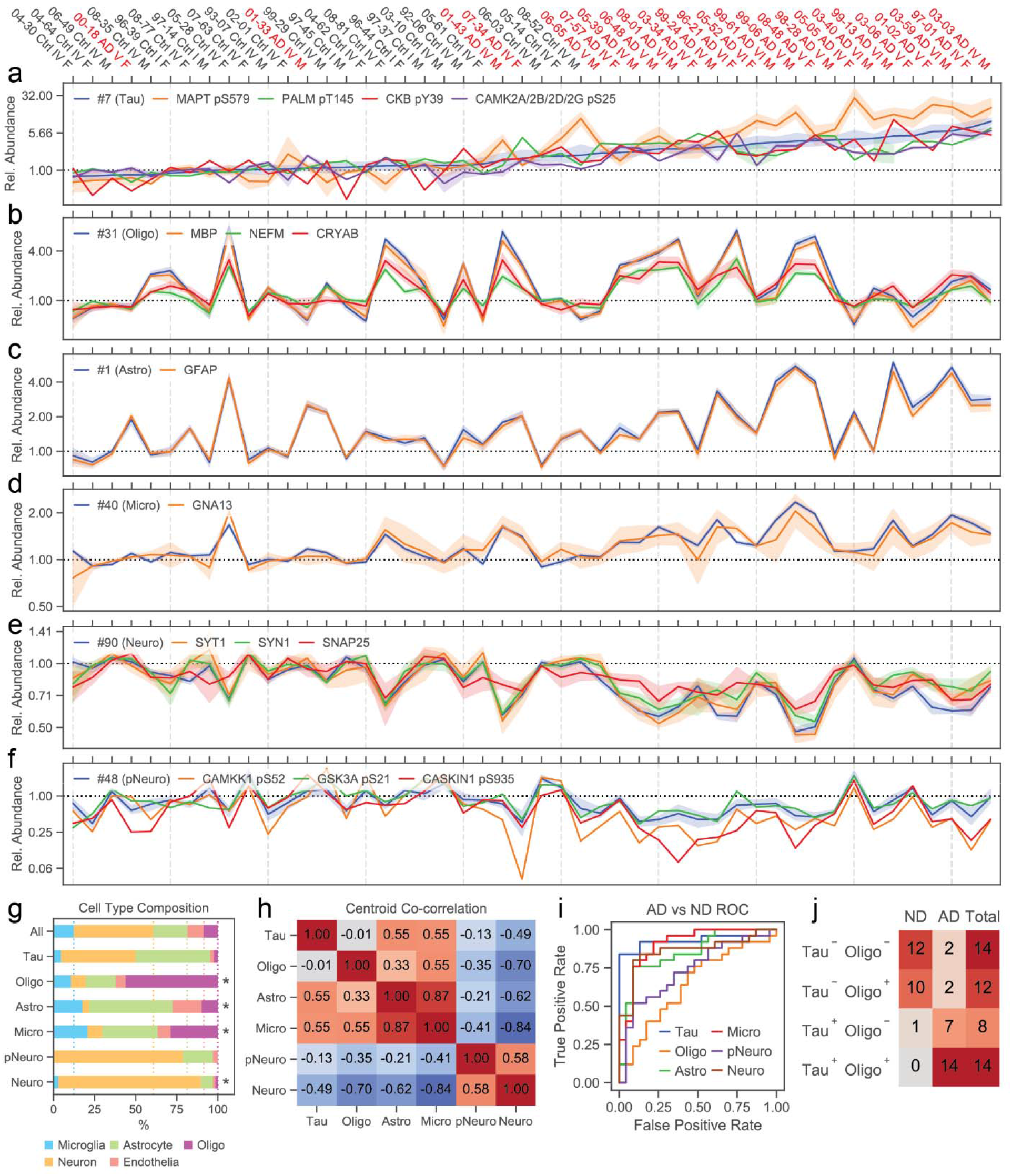
Clustering analysis identifies pathology-associated peptide modules. (a-f) Relative TMT levels for Mini-Batch K-Means cluster member peptides and selected phosphosites and proteins most correlated with each cluster centroid. Line plots show cluster centroids and member peptides for (a) Tau, (b) Oligo, (c) Astro, (d) Micro, (e) Neuro, and (f) pNeuro clusters. Line plots indicate 95% confidence intervals for all cluster peptides, all redundant phosphosites, or all peptides without PTMs (proteins). Samples for all plots are ordered by Tau centroid value shown in (a). (g) Predicted cell type composition of proteins associated with cluster centroids. Peptides seen in ≥15 patients are used for calculation. Proteins were considered if they had at least one peptide associated with cluster centroids. * indicates that cluster-associated proteins had significantly different cell type compositional make-up (χ^2^ contingency < 1e-3) compared to the entire dataset. (h) Spearman’s correlation coefficients between selected cluster centroids. (i) ROC curve predicting AD status from each named cluster centroid. (j) Patients numbers for Tau^+^ and Oligo^+^ patient stratification generated from Tau and Oligo cluster centroid values.

To confirm that these trends are not caused by tissue-sampling artifacts, we applied a series of enrichment analyses using AD-specific knowledge bases. We first examined the overlap between our dataset and a recently published large-scale proteomics analysis of AD patients^26^. We find that the Oligo protein modules in both datasets had a strong degree of overlap **(Supplementary Figure 2e)**. In addition, proteins from that study’s Astro / Micro module were enriched in our Astro and Micro clusters, while the Synapse / Neuron and Mitochondrial modules were enriched in our Neuro cluster. We next calculated the overlap between peptides associated with each cluster and the marker genes for cell type populations observed in a previous single-nuclei RNA-seq (snRNA-seq) analysis of cortical AD tissue^54^. We observed that a number of marker transcripts for the Oli0 (oligodendrocyte) population that was associated with late-stage pathology (e.g. *QDPR, CRYAB*) were similarly upregulated at the level of phosphorylation and total protein **(Supplementary Figure 2f)**. In addition, we observed that the AD-associated astrocyte (Ast1-3) and excitatory / inhibitory neuron (Ex2-8, In0-8) populations overlapped with our Astro and Neuro clusters respectively. Interestingly, the snRNA-seq analysis did not detect differential regulation of myelin proteins such as MBP, CNP, or PLP1 that were seen in both proteomics analyses **(Supplementary Figure 2g, 2h)**. Thus, we find that the cell type-specific changes that are seen in our clusters overlap with previous AD proteomics and snRNA-seq analyses^26,54^.

We next calculated the ability of our cluster centroids to estimate AD pathologies. Using a receiver operating characteristic (ROC) curve analysis, we found that Tau was the best predictor of AD status **(Figure 2i)**, in agreement with the rank order separation of AD and ND cases by Tau cluster centroid values **(Figure 2a)**. We then projected samples into 2 dimensions using principal component analysis (PCA) of the peptide matrix used for clustering. We find that AD and ND samples are separated along principle component 2 (PC2), but also that PC1 explains an even larger fraction of the data variance **(Supplementary Figure 3a)**. Samples in this space have increasing Tau centroid values along PC2 and increasing Oligo centroid values along PC1, indicating that these two clusters are closely associated with the major axes of variation **(Supplementary Figure 3b-3g)**. We further explored how other clusters related to Tau, by classifying patients as negative or positive for cluster centroid values. We selected a centroid cutoff value of 2x for Tau, and 1.25x or 0.8x for all other clusters that increased or decreased in AD patients, respectively. Based on our initial PCA analysis, we stratified patients into Tau^−^;Oligo^−^, Tau^+^;Oligo^−^, Tau^−^;Oligo^+^, and Tau^+^;Oligo^+^ groups **(Figure 2j)**. We found that this classification separated patients along PC1 and PC2 **(Supplementary Figure 3h)**. Compared to Tau;Oligo, AD and ND patients were poorly separated in PC space by either Astro;Micro or pNeuro;Neuro (**Supplementary Figure 3i-3l**). Intriguingly, we observed that several samples were Astro^+^;Micro^−^ but no samples were Astro^−^;Micro^+^ **(Supplementary Figure 3k)**. This result suggests that astrocyte activation may occur before microglial activation in the temporal progression of AD. Across the Tau;Oligo groups, Tau^+^;Oligo^+^ contained the greatest number of Astro^+^;Micro^+^ and pNeuro^+^;Neuro^+^ cases **(Supplementary Figure 3m, 3n)**. Thus, Tau is most closely linked to AD diagnosis, while Tau^+^;Oligo^+^ samples have the greatest amount of cellular pathologies across neuronal and glial cell types.

The Tau cluster may be understood in light of known AD neuropathology, but the Oligo changes are less well understood and may represent an independent stage of tissue pathology. We observe that the Oligo cluster includes neurofilament proteins NEFL, NEFM, and NEFH **(Figure 2d)** which are reported to be markers that closely track neurodegeneration^59–62^. In addition, Oligo is strongly anti-correlated with the Neuro cluster which contains SYT1 and SNAP25 **(Figure 2f, 2i)**, two synaptic markers of neurodegeneration^63,64^. This myelination/oligodendrocyte protein module has been found to be associated with the trajectory of cognitive decline in AD^28^ and has been observed in other forms of dementia caused by cerebral atherosclerosis^65^. These findings suggest that Oligo represents a general remyelination response that occurs concomitant to neurodegeneration, while Tau is most closely associated with development of AD-specific pathologies that precede neurodegeneration^66^. In this context, Tau^+^;Oligo^−^ samples are at an early stage of disease progression, while Tau^+^;Oligo^+^ samples have neurodegeneration alongside Tau pathology.

### AD pathology estimates match full proteome changes

To fully understand the molecular composition of clusters, we examined how peptides with missing quantification values were correlated with our inferred estimates of AD pathologies taken from peptide cluster centroids. We analyzed all peptides that were quantified in at least 15 samples (equivalent to ≥2 TMT10-plex analyses) with a Spearman’s rank correlation coefficient cutoff of 0.47 and p-value cutoff of 5e-3. Together, these cutoffs matched fewer than 0.23% of peptides for scrambled centroid values **(Supplementary Figure 4a-4d)**. Using these cutoffs, a total of 2609 peptides were correlated with Tau, 5554 with Oligo, 4866 with Astro, 7672 with Micro, 1863 with pNeuro, and 7200 with Neuro cluster centroids **(Supplementary Figure 4e-4g**, **Supplementary Data 2)**. This analysis identified a number of other phosphosites on MAPT that were correlated with Tau, including MAPT pT534 (2N4R isoform: pT217), which has been identified as a specific biomarker for AD^67^, as well as MAPT pT529, pS531, and pS579 (2N4R isoform: pT212, pS214, and pS262) which are associated with Tau’s seeding activity^68^ **(Supplementary Figure 4h)**. For Oligo, we identified a number of other oligodendrocyte-enriched proteins, such as CLDND1, CNDP1, and OPALIN that were closely associated with the cluster centroid **(Supplementary Figure 4i)**. Thus, we could use correlation with cluster centroids to integrate our full matrix of peptide data and find further changes reflective of cluster identity.

As there was a large degree of overlap between peptides correlated with different clusters, we decided to use partial correlation to explore the relationship of peptide changes with Tau and Oligo. We selected MAPT, MBP, and PLP1 as example proteins with peptides that increase in Tau^+^ and Oligo^+^ samples, respectively **(Supplementary Figure 5a-5d**, **Supplementary Data 4)**. We calculated the partial correlations of all peptides that were correlated with either cluster, represented in a 2D plot **(Figure 3a)**. We observed that among the correlated MAPT phosphopeptides, all were primarily correlated with Tau (Tau partial correlation MAPT pS579: ρ=0.91, p=2.7e-19; pT534: ρ=0.89, p=2.3e-17; pS713: ρ=0.92, p=1.3e-16; pS606: ρ=0.91, p=2.4e-15; **Figure 3a**). These Tau-correlated peptides were located on MAPT’s microtubule-binding domains and neighboring proline-rich domain (PNS Isoform: residues 468-685; **Supplementary Figure 5e**). In the C-terminal domain, three MAPT phosphosites (PNS Isoform: pS713, pS717, and pS721; 2N4R Isoform: pS396, pS400, pS404) were also correlated with Tau. All other peptides from the N-terminal and C-terminal domains were either uncorrelated or anti-correlated with Tau and Oligo, possibly due to degradation or modification with other PTMs. In contrast, peptides and phosphopeptides from myelin proteins (MBP, PLP1) and neurofilaments (NEFL, NEFM, NEFH) were all strongly correlated Oligo and not with Tau (Oligo partial correlation MBP: ρ=0.98, p=5.3e-31; PLP1: ρ=0.98, p=7.3e-28; NEFL: ρ=0.91, p=3.0e-19; NEFM: ρ=0.94, p=2.0e-22; NEFH: ρ=0.91, p=4.2e-16; **Figure 3b, 3c**, **Supplementary Figure 5c, 5d)**. The one exception was a peptide mapping to MBP_31-37_ that was anticorrelated with Oligo **(Supplementary Figure 5f)**. This peptide is specific for the Golli MBP isoforms that are only expressed during development^69^. All other MBP peptides were correlated with Oligo and mapped to positions after the 134 residue start site for MBP isoforms 3-6. We finally examined the proteome, estimating protein abundances from median peptide levels and calculating their predicted cell types. We observed that oligodendrocyte proteins were primarily correlated with Oligo, neuronal proteins were primarily anticorrelated with Oligo, and microglia and astrocyte proteins were partially correlated with both Tau and Oligo to varying degrees **(Figure 3d)**. The anticorrelated neuronal proteins included synaptic markers of neurodegeneration^63,64^ (Oligo partial correlation SYT1: ρ=−0.86, p=4.7e-15; SNAP25: ρ=−0.62, p=2.9e-6). These results match the patterns of cell-type enrichment and correlation between cluster centroids that we had previously observed **(Figure 2g, 2h)**. Thus, our expanded correlation preserves the cell-type association established with a smaller data set quantified across all patients.

**Figure 3.**
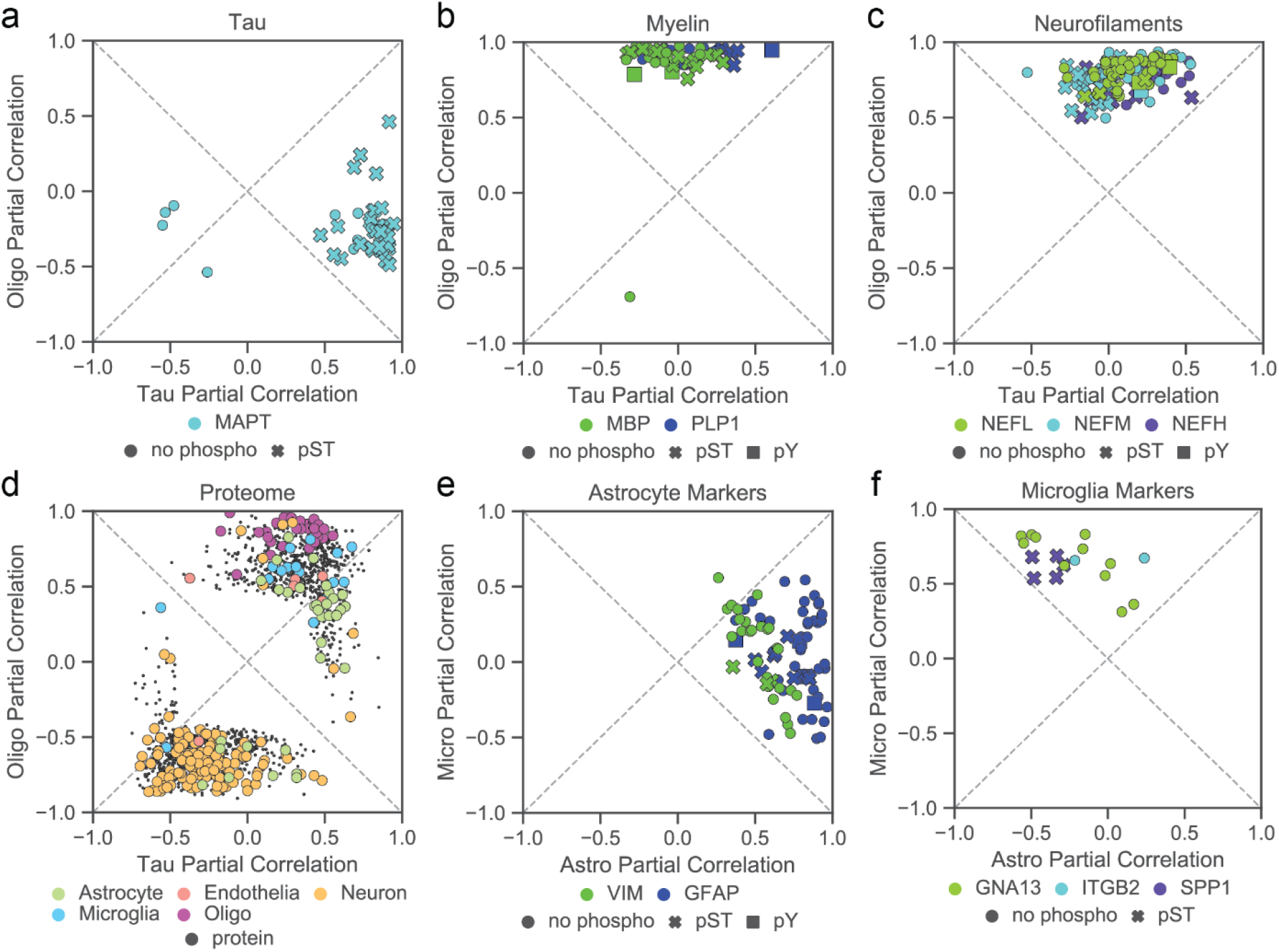
Correlation analysis finds pathology-associated change in the full peptidome. (a-c) Partial correlation with Tau and Oligo cluster centroids for (a) MAPT peptides, (b) myelin proteins, and (c) neurofilament proteins. Only peptides that are seen in ≥15 patients and significantly correlated with either Tau or Oligo are shown. Points are colored by protein ID and stylized by the presence of PTMs. Dotted lines on the diagonal are shown as reference for relative correlation. (d) Partial correlation with Tau and Oligo for all protein levels estimated from all non-phospho peptides. Points are colored by proteins’ predicted cell type. (e-f) Partial correlation with Astro and Micro for selected (e) astrocyte-enriched and (f) microglia-enriched peptides.

We applied a similar methodology to see if we could identify other microglial and astrocytic activation markers that were correlated with their respective clusters. We find that the astrocyte activation markers GFAP and VIM are strongly correlated with Astro at the level of phospho and non-phospho peptides (Astro partial correlation GFAP: ρ=0.97, p=1.4e-28; VIM: ρ=0.77, p=1.6e-10; **Figure 2e**, **Supplementary Figure 5g, 5h)**. In contrast, the microglia-enriched protein GNA13 and activation markers AIF1 (IBA1), ITGB2 (CD11B), and SPP1 were all more strongly correlated with Micro than Astro (Micro partial correlation GNA13: ρ=0.83, p=2.7e-13; SPP1 pS191, pS195: ρ=0.83, p =7.4e-6; ITGB2: ρ=0.67, p=9.0e-5; AIF1: ρ=0.58, p=7.9e-3; **Figure 2f**, **Supplementary Figure 5i-5l)**. These results are notable, as previous large-scale proteomics analyses have not been able to decouple astrocyte from microglial changes^26^.

### AD pathologies are correlated with dysregulated kinase networks

We next examined how phosphorylation sites on different kinase, phosphatase, and receptor signaling factors were correlated with our cluster centroids. Out of all phosphosites on these signaling factors that were correlated with Tau, Oligo, Astro, or Micro, 85% were correlated with Tau or Oligo **(Figure 4a)**. To understand how known AD signaling factors were correlated with these clusters, we selected a subset of known MAPT kinases, including BRSK1, BRSK2, TNIK, and TTBK1, and plotted their correlation with Tau and Oligo centroids **(Figure 4b)**. We observed several phosphosites on TNIK, BRSK1, and BRSK2 that were primarily correlated with Tau, matching their upregulation in Tau^+^ patient groups (TNIK pS680: FC=1.92, p=2.44e-4; BRSK1 pS443: FC=1.60, p= 4.40e-6; BRSK2 pS423: FC=1.97, p=6.12e-7; **Supplementary Figure 6a-6c**, **Supplementary Data 4**). In contrast, TTBK1 phosphosites were located closer to the diagonal of the plot, reflecting that it was phosphorylated to a larger degree in Tau^+^;Oligo^+^ samples (TTBK1 pS483: FC=2.52, p=5.43e-6; **Supplementary Figure 6d**). The presence of phosphosites on these kinases that are highly correlated with pathology is striking as we did not observe a corresponding change in any of the non-phospho peptides, suggesting an increase in the stoichiometric ratio of phosphorylation on those sites **(Supplementary Figure 6e-6h)**. We applied a similar analysis to the several cyclin-dependent kinases (CDKs) and observed that phosphosites on CDK5, CDKL5, CDK16, CDK17, and CDK18 were correlated with both Tau and Oligo **(Figure 4c)** and significantly upregulated in Tau^+^;Oligo^+^ samples (CDK5 pY15 FC=1.8, p=1.2e-2; CDKL5 pY171: FC=3.1, p=2.5e-5; CDK16 pS153: FC=3.4, p=2.9e-6; CDK17 pS180: FC=2.9, p=5.0e-10; CDK18 pS132: FC=3.6, p=2.7e-9; **Supplementary Figure 6i-6m**, **Supplementary Data 4**). For CDK5 and CDKL5, we observed that non-phosphopeptides were significantly downregulated in this group, while only CDK18 had peptides that were also upregulated (CDK5 FC=0.86, p=5.0-5; CDKL5: FC=0.67, p=2.7e-3; CDK18: FC=1.5, p=9.0e-3; **Supplementary Figure 6n-6r**). Among these phosphosites, CDK5 pY15 potentiates kinase activation through Aβ_42_-c-Abl signaling^70,71^; CDKL5 pY171 is an autophosphorylation site and its mutation abolishes enzymatic function^72^; CDK16 pS153 is a Protein Kinase A (PKA) substrate that blocks the protein’s interaction with CCNY and inhibits its function^73^; CDK17 pS180 is elevated in Aβ-expressing cell lines^74^. While CDK5 and CDK18 are known to have altered signaling in the AD brain^75,76^, this observation has not been previously made of CDKL5, CDK16, and CDK17.

**Figure 4.**
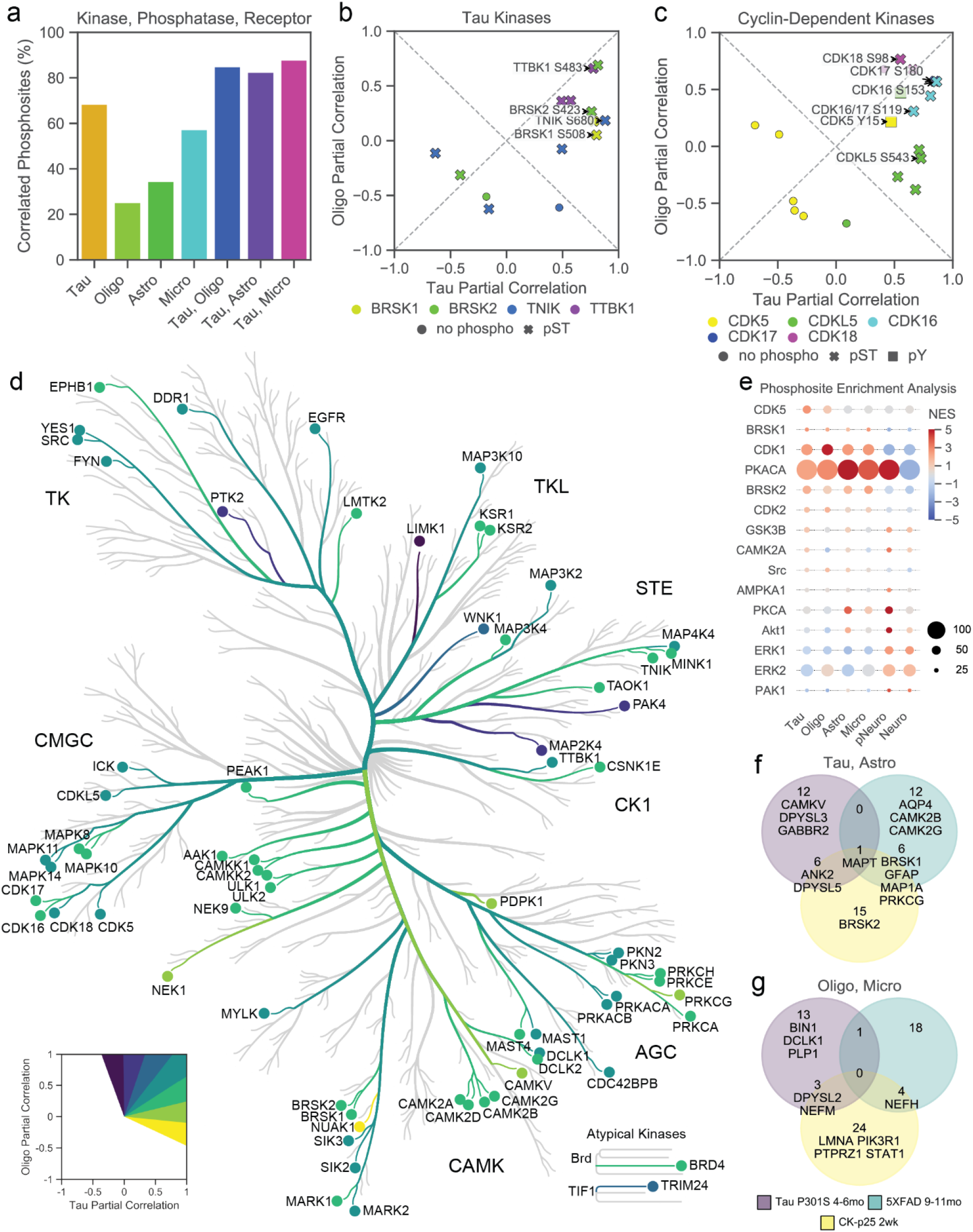
Kinome map of the pathology-associated phosphoproteome. (a) Percentage of kinase, phosphatase, and receptor phosphosites that are significantly correlated with Tau, Oligo, Astro, or Micro. (b-c) Partial correlation with Tau and Oligo for selected (b) Tau kinase and (c) cyclin-dependent kinases. (d) Kinome map of all phosphopeptides that were significantly associated with Tau or Oligo cluster centroids. Branches and nodes are colored by partial correlation with Tau and Oligo as shown on inset panel. (e) Phosphosite enrichment analysis applied to the phosphoproteome dataset for each cluster centroid. Normalized-enrichment scores are shown for phosphosite sets that were significantly associated with at least one cluster centroid. Circle sizes correspond to the number hits for each phosphosite set. (f-g) Phosphoproteins that were identified to have at least one shared phosphosite with three mouse models of neurodegeneration. Proteins are shown as a Venn diagram between upregulated phosphosites and (f) Tau or Astro and (d) Micro or Oligo cluster centroids. Numbers indicate the total number of phosphoproteins in each overlapping set and selected protein names are shown.

To aid visualization, we built a map of all kinases with phosphosites that were significantly correlated with Tau or Oligo variables **(Figure 4d)**. We categorized nodes based on their relative correlation with Tau and Oligo. This approach highlighted proteins from all kinase families within the human kinome that were associated with Tau and Oligo. Out of the 63% of these kinases that had protein quantification, only 23% were significantly upregulated at a protein level in Tau^+^ or Oligo^+^ samples, while 46% were downregulated in these groups **(Supplementary Figure 7a**, **Supplementary Data 4**). The associated kinases included MAPK8/10 (pY185/pY223: FC=2.4, p=1.3e-4), MAPK11 (pY182: FC=4.7, p=1.8e-5), MAPK14 (pY182: FC=2.5, p=1.5e-5), Protein Kinase A C-subunit α/β (PKACA/B pS140: FC=1.8, p=2.7e-5), Protein Kinase C α (PKCA pT228: FC=1.5, p=1.9e-3), and CAMKII α/β/δ/γ (pS25/pS26: FC=3.3, p=2.9e-6; **Supplementary Figure 7b-7h**), which have previously found to be activated in AD^7,8,77–79^. Among the phosphosites that we identified, MAPK8 pY185 / MAPK10 pY223 increase kinase activity and mediate a stress response^80^. Similarly, MAPK14 pY182 is necessary for kinase activity and mediates a stress response^81^ and MAPK11 pY182 has a high degree of homology to MAPK14 that likely confers similar functional characteristics. PKACA/B are subunits of another kinase that phosphorylates Tau^7^, and pS54 and pS140 are both autophosphorylation sites on these proteins^82^. PKCA is a kinase that mediates synaptic pathologies caused by Aβ^8^; PKCA pT228 is phosphorylated by MST2 and is necessary for its activation in mice^83^. For CAMKII, we observe several phosphosites on its protein subunits that are significantly upregulated in Tau^+^;Oligo^+^ samples, including pY17, pS25, pS71, pS145, pS257, and pS276, even as total protein levels of CAMKII subunits are significantly downregulate in this same group **(Supplementary Figure 7h**, **Supplementary Data 4)**. While most of these phosphosites are of unknown function, pT306 has been reported to block CAMKII’s interaction with Calmodulin, inhibiting its kinase activity^84^. Additionally, CAMKII’s pT286 autophosphorylation site^85^ is downregulated in this group and correlated with pNeuro, indicating that CAMKII has reduced total activity in samples with neuronal pathology **(Supplementary Figure 7h**, **Supplementary Data 2)**. These findings point to a complex role of phosphorylation PTMs in regulating CAMKII activity in AD.

We next examined whether other kinase associations represented novel insight. Within the tyrosine kinase family tree, SRC-family kinases (SFKs) act as key signaling nodes in mediating tyrosine receptor signaling^86^. We observed increased phosphorylation on SRC in Tau^+^;Oligo^+^ samples (pS75: FC=1.5, p=1.2e-3; pY187: FC=1.4, p=6.0e-5; pY439*: FC=2.7, p=1.2e-3; **Supplementary Figure 7i**). SRC pS75 has been identified as a substrate of CDK5 that inhibits SRC function^87^. While the exact function of SRC pY187 is not currently known, the site is on SRC’s SH2 domain and bears strong sequence homology to similar autoinhibitory sites on other SFKs^88^. Alongside SRC pS75, we see that LMTK2, another CDK5 substrate^89^, has increased phosphorylation in Tau^+^;Oligo^+^ samples (pS1450: FC=1.6, p=6.4e-3; **Supplementary Figure 7j**). Downstream of SRC, we observe that PTK2 has increased phosphorylation in this same group (pY925: FC=1.6, p=2.9e-4; **Supplementary Figure 7k**). This phosphosite is a SRC-specific substrate that is necessary for PTK2’s regulation of cell motility in human cell lines^90^. Among the other tyrosine kinases, we observed that DDR1 has increased phosphorylation in both Tau^+^ and Oligo^+^ samples (pY796: FC=2.1, p=3.1e-4; pY792: FC=1.6, p=5.5e-3; **Supplementary Figure 7a, 7l**). These two phosphosites are on the conserved activation loop of DDR1^91^ and inhibition of this receptor has recently been shown to decrease inflammation in mouse models of AD and Parkinson’s disease^92–94^. We also observed that an EGFR autophosphorylation site^33^ is increased in Tau^+^;Oligo^+^ samples (pY1175: FC=2.9, p=9.4e-4; **Supplementary Figure 7m**). In total, we find that our collection of Tau- and Oligo-correlated phosphosites includes a number signaling factors that are known to be active in AD as well as novel associations on sites linked to increased kinase activity.

### AD kinase networks overlap with substrate and mouse model phosphoproteomes

To test whether the phosphosites downstream of the pathology-associated kinases were also dysregulated, we applied phosphosite enrichment analysis (PSEA) using publicly available kinase-substrate datasets^56^. By calculating enrichment scores for different cluster correlates, we find that phosphosites regulated by BRSK1, BRSK2, CDK5, PKACA, CDK1, CAMK2A, PKCA, GSK3B, ERK1, and ERK2 are associated with one or more cluster centroids **(Figure 4e)**. These findings are supported by the direct association of phosphosites on BRSK1 and BRSK2 **(Supplementary Figure 6f, 6g)**, CDK5 **(Supplementary Figure 6n)**, CAMK2A **(Supplementary Figure 7a)**, PRKCA **(Supplementary Figure 7a)**, GSK3A/B **(Figure 2f**, **Supplementary Data 2)**, and MAPK3 and MAPK1 (ERK1 and ERK2; Supplementary Data 2) with the most strongly enriched clusters. In the case of PKACA, we find that some downstream phosphosites increase with Tau, Astro, and Oligo, while others decrease with pNeuro **(Supplementary Figure 8a-8d)**. Among the Tau- and pNeuro-correlated PKACA substrates, CAMKK1 pS475 and pS52 are increased and suppressed, respectively, in the presence of Ca^2+^/Calmodulin^95^. Many of the Astro- and Oligo- correlated PKACA phosphosites are located on astrocyte- and oligodendrocyte-enriched proteins, suggesting that this kinase also dysregulated these cell types as well. Altogether, we find that kinase phosphosites are correlated with substrate phosphopeptides changes and reflect signaling pathways known to be activated in AD. Our data strengthens support for these factors and adds a new layer with several novel kinase associations.

We finally examined the overlap between the phosphoproteomes of AD and three mouse models of neurodegeneration^22^. In this previously collected dataset, we analyzed the phosphoproteome of CK-p25, 5XFAD, and Tau P301S mouse brain tissue using a similar methodology to this study. Out of all of the differentially phosphorylated sites in mice, 96.7% could be mapped by homology to human proteins **(Supplementary Data 5)**. We observed that the CK-p25 mouse model with two weeks of p25/Cdk5 activation had the greatest number of overlapping phosphosites that shared directional changes with AD pathologies **(Supplementary Figure 8e)**. Interestingly, MAPT emerged as the one protein with phosphorylation sites that were associated with pathology in each mouse model and in human patients **(Figure 4f-4g**, **Supplementary Figure 8f)**. We observed a number of aforementioned signaling factors that also had phosphosites enriched in one or two mouse models, including BRSK1, BRSK2, CAMKII, CAMKV, and PTPRZ1. This comparison also allows us to tie phosphorylation on these signaling factors to pathologies driven by p25/Cdk5, Aβ, or phospho-MAPT activation in each respective mouse model. These results may inform the future choice of mouse models to study particular signaling pathway dysregulation events.

### Statistical modeling links the phosphoproteome to AD histopathologies

To better understand the associations between peptide changes and histopathology variables, we generated a series of partial least squares regression (PLSR) models. We used PLSR to link our cluster centroids to each component of the full matrix of histopathology measurements **(Figure 5a**, **Supplementary Data 1)**. We first aimed to reduce the size of the histopathology matrix to variables that were relevant to the cortex tissue that we analyzed. By building separate PLSR models with K-fold cross validation analysis for each outcome in the histopathology matrix, we determined that a subset of pathology variables could be predicted by PLSR from peptide data with moderate accuracy (*e.g.* Consortium to Establish a Registry for Alzheimer Disease neuropsychological battery (CERAD-NP, NIA-Reagan (NIA-R), plaque score, and temporal tangles (TangleT); Q^2^ ≅ 0.39 - 0.60) **(Figure 5b)**. Although our models do not have predictive accuracy for AD risk variables such as APOE genotype or White Matter scores, our patient cohort did not have any cases of the most severe APOE ε4/ε4 alleles and there are few cases with White Matter score > 0 in the temporal lobe where we sampled tissues.

**Figure 5.**
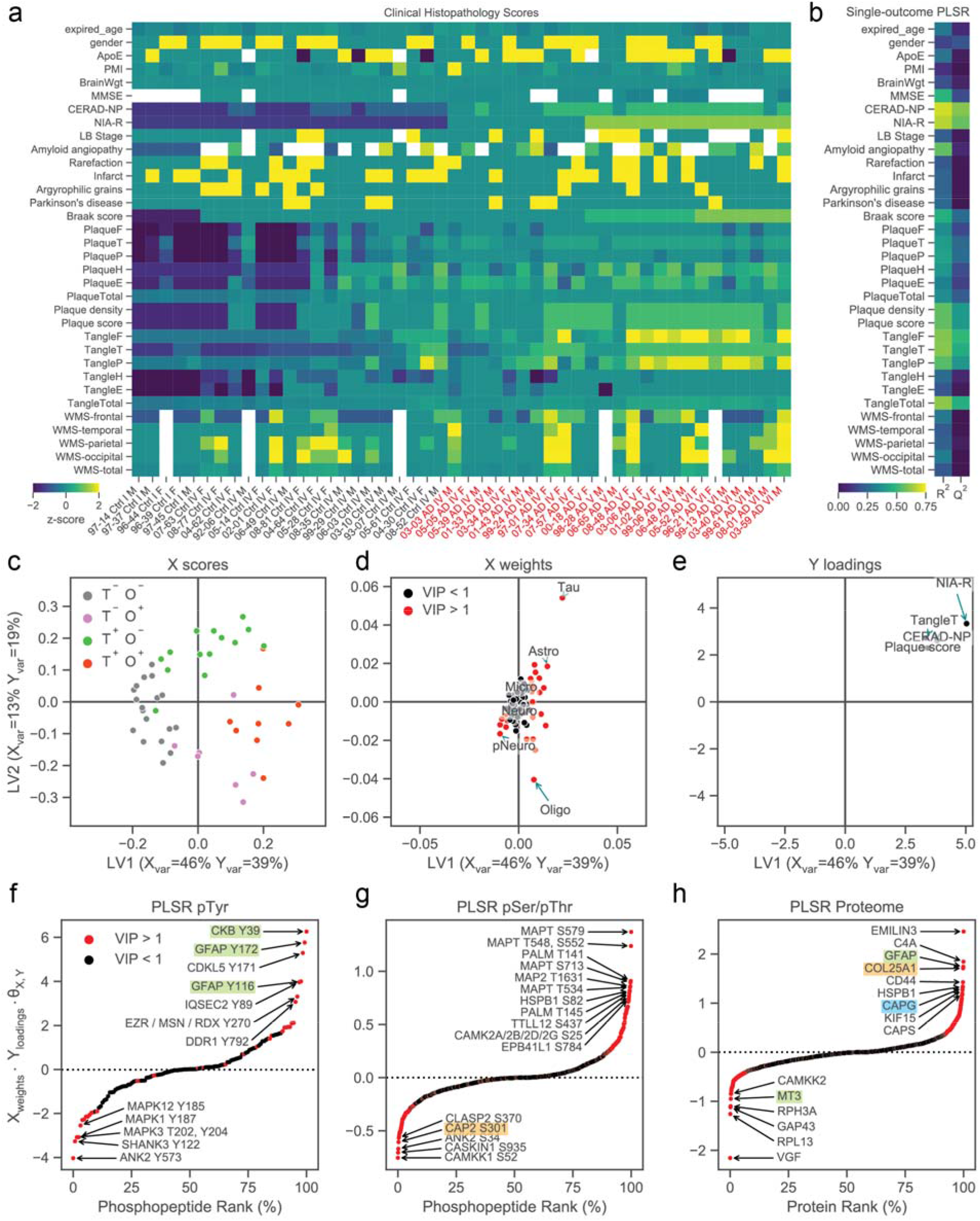
PLSR model ties clusters and peptides to AD histopathologies. (a) Clinical histopathology measurements analyzed by PLSR for each patient. Rows are z-score normalized and missing values are shown as white. (b) K-fold validation of single-outcome PLSR models for each histopathology variable. R^2^ and Q^2^ values are shown for training and test data predictions respectively. (c-e) Multivariate PLSR model using cluster centroids and select clinical histopathology variables. (c) X scores, labeled by Tau (T) and Oligo (O) status, (d) X weights, and (e) Y loadings vectors are shown in latent variable (LV) space. The percent variance explained by the first two components of the model for the X and Y matrices are shown on the axis labels. (f-h) Peptide-pathology association scores generated from peptide-centric multivariate PLSR models built from (f) pTyr peptides, (g) pSer/pThr peptides, and (h) estimated protein abundances.

In order to interpret the covarying relationship between peptide cluster centroids and pathology outcomes, we generated a PLSR model linking cluster centroids to a multivariate Y block matrix composed of the non-redundant predictable pathology variables (CERAD-NP, NIA-R, plaque score, and TangleT). We found that a 2-component PLSR model explained ~57% of the data variance for both X and Y matrices **(Figure 5c)**. We then plotted this model’s X scores, X weights, and Y loadings vectors in latent variable (LV) space to visualize how centroid and pathology measurements covary. Samples were separated by their X scores values on LV1 based on Tau status and LV2 based on Oligo status **(Figure 5c)**. We examined the X weight vectors for this model and found that the Tau, Astro, Oligo, and pNeuro centroids all strongly contributed to the final model (VIP score > 1; **Figure 5d**). In LV space, Tau, Astro, and pNeuro weight vectors strongly contributed to LV1 separation, while Oligo was the strongest contributor to LV2 separation **(Figure 5d)**. Furthermore, the Tau/Astro and pNeuro clusters have roughly opposite directions from the origin **(Figure 5d)**, in agreement with these groups being upregulated and downregulated respectively in AD patients **(Figure 2j**, **Supplementary Figure 3k, 3l)**. Finally, we examined the Y loadings vectors for this model and found that plaque score, TangleT, NIA-Reagan, and CERAD-NP all occupy the same LV space **(Figure 5e**, **Supplementary Figure 9a**-**8c)**. We tested the accuracy of this multivariate PLSR model and found that it had a similar predictive accuracy to the single-outcome models **(Supplementary Figure 9d)**. Overall, our regression analysis allows us to relate our cluster centroids to patient AD histopathology measurements. Intriguingly, there were not any histopathology variables that were strongly associated with the negative direction of Oligo changes on LV2, yet AD patients were well distributed along the LV2 axis between Tau^+^ (high LV2) and Oligo^+^ (low LV2). These results highlight the need for post-mortem characterization of oligodendrocyte pathology alongside other traditional AD pathologies.

To understand which individual peptides were most predictive of AD pathologies, we built a series of peptide-centric multivariate PLSR models. We find that peptide-centric models give similar patient-histopathology association scores as the model built on cluster centroids **(Supplementary Figure 9e)**. To visualize the relation between peptides and histopathologies, we rank ordered peptides by their association with AD pathologies in LV space **(Supplementary Data 6)**. We first analyzed a model generated from pTyr peptides measured in all samples and found that creatine kinase B-type (CKB) pY39 was the most strongly associated pTyr peptide with pathology **(Figure 5f)**. Focal creatine deposits have been observed in AD brains^96^, and CKB is reported to have post-translation oxidation and downregulated activity in AD^97^. Creatine and cyclocreatine have been shown to have some positive effects in mouse models of neurodegeneration and have been suggested as a dietary supplements to offset the loss of CKB activity^98–100^. Alongside CKB pY39, we found several other phosphosites on CKB that are increased in Tau^+^ samples even as total CKB protein does not change **(Supplementary Figure 9f)**. These phosphosites may contribute to CKB’s reduced activity, although their exact function is not yet known. IQSEC2, a component of the NMDA receptor complex which is overactivated by Aβ in AD^101,102^, was also associated with pathology. Phosphosites on CDKL5 and DDR1 were also associated with AD (**Figure 5f**), strengthening our findings from kinase correlation analysis. In the other direction, we saw that phosphosites on the synaptic proteins ANK2 and SHANK3 as well as phosphosites on the activation loops of MAPK1 and MAPK3 were negatively associated with AD **(Figure 5f)**. This finding matches the prediction from PSEA that ERK1 and ERK2 substrates were associated with pNeuro and Neuro **(Figure 3i)**. This contrasts with MAPK1 and MAPK3 being significantly upregulated at the level of total protein in Tau^+^;Oligo^+^ samples **(Supplementary Data 4)**, suggesting that ERKs are less activate at a bulk tissue level in AD. The upregulation of ERK proteins may be a compensatory mechanism or due to differences in ERK regulation in different cell types in the brain. While acute Aβ stimulation has been shown to activate the ERKs in neurons, the total level of phosphorylation on the activation loops of ERK is highly variable in AD tissues^103^ and had not be previously associated with specific disease processes.

In a second PLSR model built from pSer/pThr peptides, we found that phosphosites on MAPT were among those most strongly associated with AD pathologies **(Figure 5g)**. These sites include the aforementioned MAPT pS579, pS713, and pT534 phosphosites that were strongly correlated with the Tau centroid. We also observed that Paralemmin-1 (PALM) pT141 and pT145 are strongly associated with AD **(Figure 5g)**. PALM is a modulator of plasma membrane dynamics^104^ with an uncharacterized role in AD. In the other direction, the pNeuro cluster peptides CAMKK1 pS52 and CASKIN1 pS935 were among the most negatively-associated phosphopeptides in this model **(Figure 5g)**. Compared to the conventional synaptic markers SYN1 and SYT1 which are significantly downregulated only in Neuro^+^ samples, these phosphosites were primarily downregulated in pNeuro^+^ samples **(Supplementary Figure 9g-9j)**. These findings indicate that the neuronal synaptic phosphoproteome is affected in AD, even in the absence of total synaptic protein downregulation.

In a final PLSR model built on the proteome, we find that the secreted glycoprotein EMILIN3 and complement protein C4A were the proteins most strongly associated with pathology **(Figure 5h)**. Of these two proteins, EMILIN3 was specifically upregulated only in Tau^+^ samples **(Supplementary Figure 9k-9l)**. Little is known about EMILIN3 in the context of the brain, and, to our knowledge, this is the first report of its association with AD. VGF, a marker reported to be downregulated in AD^28,105^, was the most negatively-associated protein with pathology in this model and was significantly downregulated only in Tau^+^ samples **(Supplementary Figure 9m)**. ROC analysis of the proteins and phosphosites highlighted by our PLSR models finds they are each predictive of AD status **(Supplementary Figure 9n)**. Overall, our analysis recapitulates known protein and phosphopeptide links to AD and reveals new markers and signaling factors, such as the kinases CKB, CDKL5, and DDR1, that are closely associated with disease.

## Discussion

In this study, we used phosphoproteomics to explore cellular signaling network changes in AD. We followed a data collection strategy that enabled the quantification of low-abundance pTyr phosphopeptides alongside the pSer/pThr phosphoproteome and proteome from 48 AD and age-matched control patients. By capturing a deep molecular profile of the phosphoproteome alongside the proteome for individual patient samples, we were able to build a map of how signaling factors become activated alongside other pathological processes. Our analysis uncovered co-correlated sets of peptides including: (1) “Tau,” a cluster associated with Tau pathology and (2) “Oligo,” a cluster containing oligodendrocyte, myelination, and neurofilament proteins. Tau was closely associated with AD diagnosis, while Oligo was more closely associated other cellular pathology peptide modules. In addition, we identified neuronal phosphopeptide and protein modules that were negatively associated with Tau and oligodendrocyte pathologies. We observed phosphosites on several AD-associated kinases that were correlated with pathological changes, as well as kinase-substrate networks that followed similar trends. We additionally found several novel kinase phosphosites that were correlated with these pathologies, including CDKL5, CDK16, and CDK17, as well as tyrosine kinases SRC, EGFR, DDR1, and PTK2. Our dataset adds a new layer of understanding on how these signaling factors correlate with early- and late-stage AD pathologies.

Although our analysis was of bulk tissue, many of the differentially expressed proteins were found to be primarily expressed in glial cells. Among the previously reported markers of neurodegeneration^60–64,106^, we observed that the Oligo cluster is strongly correlated (Spearman ρ > 0.9) with neurofilament proteins and strongly anticorrelated (Spearman ρ < −0.7) with synaptic proteins. One major question arising from our work is the nature of this myelination response. Previous proteomics analyses have linked myelination to cerebral atherosclerosis as well as the trajectory of cognitive decline in AD^26,28,65^. In addition, a subset of these proteins were found to be upregulated at a transcript level in single-nuclei RNA-seq analyses of AD oligodendrocytes^54^. A previous analysis of a mouse model of tauopathy has found that oligodendrogenesis increases in hippocampal grey and white matter regions prior to locomotive and memory impairments^107^. We have similarly observed increased phosphorylation on PLP1 by 6 months of age in the Tau P301S model. However, our AD proteomics analysis also finds a number of Oligo^+^ patients without Tau pathology. This suggests that myelination occurs as a general response concomitant to neurodegeneration downstream of Tau pathology alongside other neurotoxic insults.

Overall, our analysis highlights the importance of integrating the phosphoproteome with proteome and transcriptome datasets. While our proteome dataset captured changes in oligodendrocyte and neuronal protein abundances, the pSer/pThr and pTyr phosphoproteomes were necessary to directly measure Tau pathology as well as a downregulated neuronal phosphopeptide module. The phosphoproteome also provides direct measurement of the regulatory sites on protein kinases and other signaling factors. We identified a number of proteins that are associated with AD at the level of phosphorylation even as the proteins themselves are unchanged or downregulated. These phosphorylation changes were most likely to be associated with Tau pathology, supporting the hypothesis that protein phosphorylation events mediate signaling events that are pre-emptive of neurodegeneration. Our dataset allows us to separate signal events associated with early-vs. late-stage AD pathologies as well as other forms of neurodegeneration. By targeting the signaling events occurring at an early-stage of AD, we may be able to delay or prevent disease progression.

## Acknowledgements

We thank members of the White, Tsai, and Lauffenburger laboratories for numerous discussions and feedback. N.M. was partially supported by the NIH Biotechnology Training Grant T32GM008334. M.L. was partially supported through the US Army Research Office Cooperative Agreement W911NF-19-2-0026 for the Institute for Collaborative Biotechnologies and the National Science Foundation Graduate Research Fellowship Program. We are grateful to the Banner Sun Health Research Institute Brain and Body Donation Program of Sun City, Arizona for the provision of human brain tissue. The Brain and Body Donation Program has been supported by the National Institute of Neurological Disorders and Stroke (U24 NS072026 National Brain and Tissue Resource for Parkinson’s Disease and Related Disorders), the National Institute on Aging (P30 AG19610 Arizona Alzheimer’s Disease Core Center), the Arizona Department of Health Services (contract 211002, Arizona Alzheimer’s Research Center), the Arizona Biomedical Research Commission (contracts 4001, 0011, 05-901 and 1001 to the Arizona Parkinson’s Disease Consortium) and the Michael J. Fox Foundation for Parkinson’s Research.

## Author Contributions

N.M. and F.W. designed the study. D.M. collected human brain tissue samples. N.M. and F.H.R. collected proteomics data sets. N.M. wrote proteomics data integration and enrichment software. N.M. and M.L. performed statistical analyses. N.M., M.L., and F.M.W. wrote the manuscript.

## Materials & Correspondence

Correspondence and requests for materials should be addressed to F.M.W.

## Supplementary Information

Supplementary Data 1. **Demographic information and clinical histopathology measurements for individual patients**.

Supplementary Data 2. **Proteome and phosphoproteome cluster correlations**. Table includes the spearman’s correlation and associated p-values between each peptide and each cluster centroid.

Supplementary Data 3. **Peptides cluster identities**. Cluster identities and centroid values for peptides quantified in all samples.

Supplementary Data 4. **Proteome and phosphoproteome cluster fold changes**. Table includes fold changes and p-values for each peptide and cluster group against their respective negative cases.

Supplementary Data 5. **Mouse phosphoproteome overlap**. AD mouse model phosphosites from Morshed et al. (2020) that had homologous phosphosites detected in the human AD phosphoproteome.

Supplementary Data 6. **Peptide scores and VIP values for peptide-centric PLSR models**. Table includes tabs for PLSR models built from pTyr, pSer/pThr, Protein, and all data.

---

**Supplementary Figure 1.**
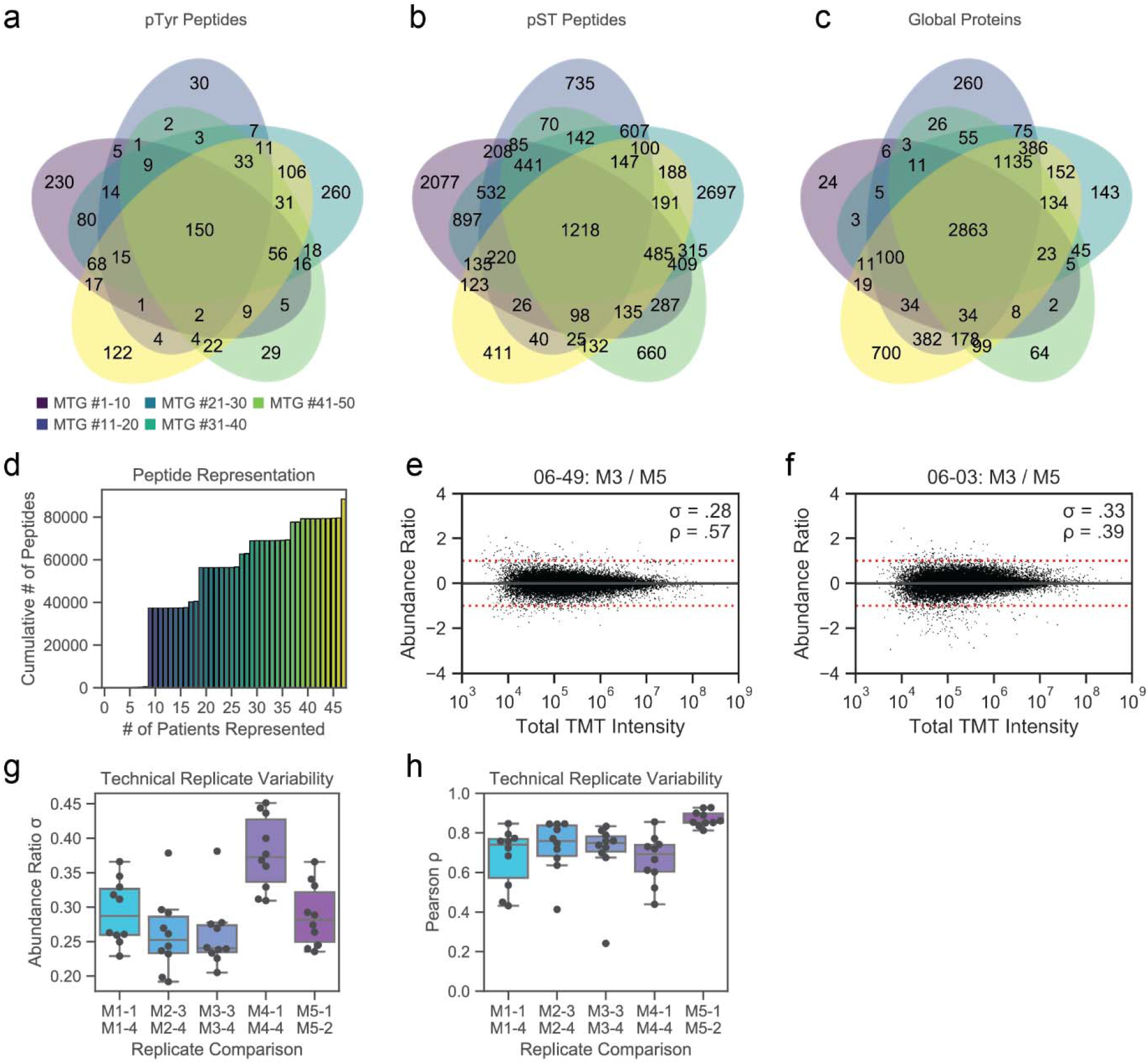
Proteome and phosphoproteome dataset statistics and quantification reproducibility. (a-c) Venn diagram showing peptide overlap between (a) pTyr sites, (b) pSer/pThr sites, and (c) proteins identified in each 10-plex analysis. (d) Cumulative number of peptides identified compared to the number of patients with peptide quantification. (e-f) Reproducibility analysis of peptide quantification after data integration and normalization. Show are the log_2_ quantification ratios between two separate brain samples from patients (e) 06-49 and (f) 06-03. Peptides are separated on the x-axis by log_10_ total TMT intensity from all PSMs in both 10-plex analyses. The standard deviation of abundance ratios and Pearson’s correlation coefficient between all peptide replicates are listed in the panels. (g-h) Reproducibility of peptide quantification between technical replicates for each TMT10-plex analysis. Shown are (g) the standard deviation of abundance ratios and (h) the Pearson’s correlation coefficient calculated over all peptides.

**Supplementary Figure 2.**
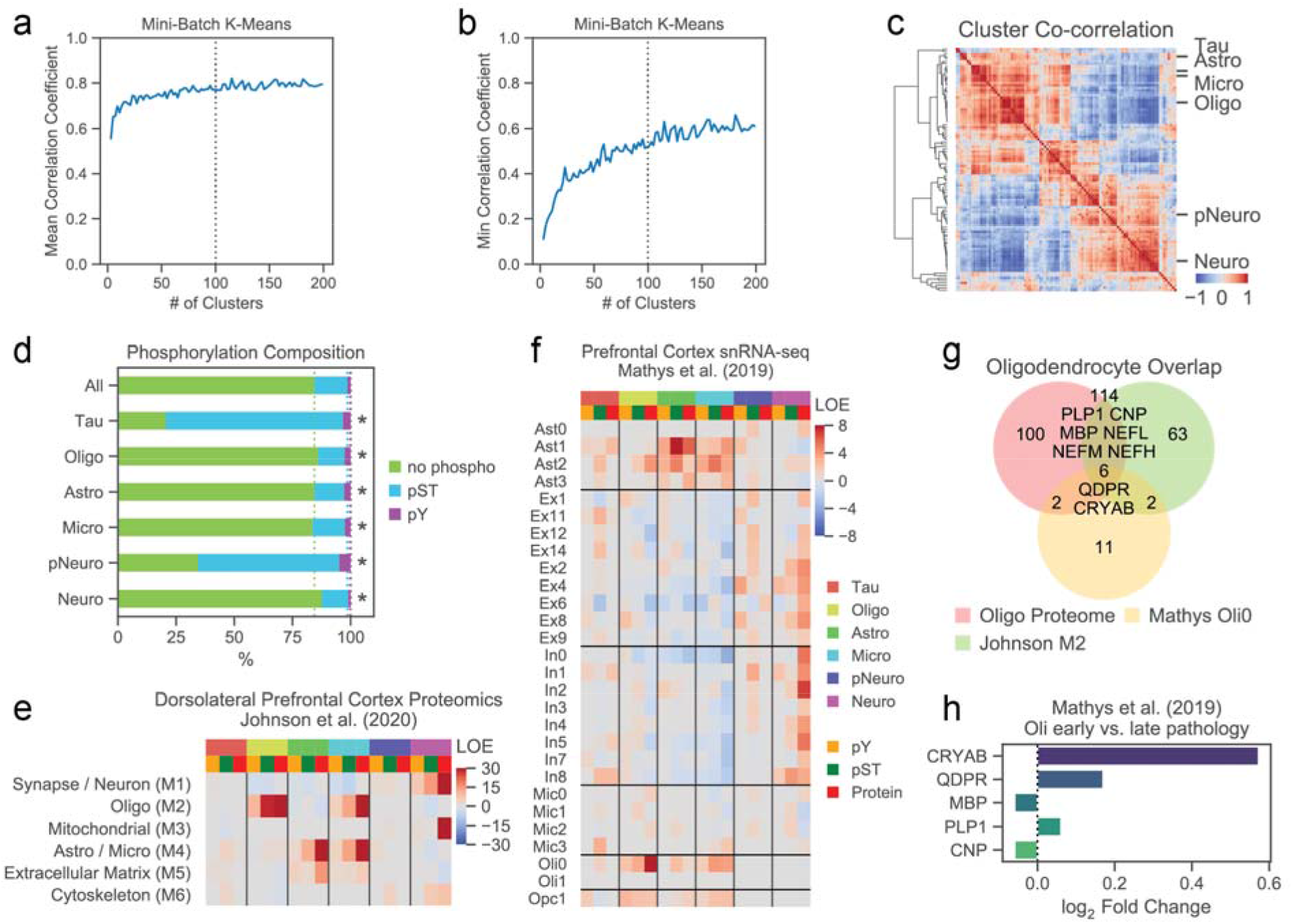
Accessory clustering and dataset overlap analyses. (a-b) Properties of clusters generated by clustering with increasing number of clusters. Shown are the (a) mean Pearson correlation coefficient for cluster peptides with their respective cluster centroid, averaged over all clusters, and (b) minimum absolute Pearson correlation coefficient for cluster peptides with their respective cluster centroid, summed and normalized over all clusters. Dotted line indicates the final cluster count used for downstream analysis. (c) Co-correlation heatmap between all cluster centroids, group by cluster. Selected named clusters are shown on the y-axis. (d) Phosphopeptide composition of all peptides associated with cluster centroids. Peptides are labeled as ‘pY’ if they contain at least one pTyr site, ‘pST’ if they contain at least one pSer or pThr site, or ‘no phospho’ if they are not phosphorylated. Statistics calculations are same as **Figure 2g**. (e) Binomial enrichment for proteome module marker proteins identified in Johnson et al. (2020). Heatmap shows log-odds enrichment (LOE) values for the overlap of marker transcripts with pTyr, pSer/pThr, and proteome datasets that are correlated with cluster centroids. (f) Binomial enrichment for single-nuclei populations markers identified in Mathys et al. (2019). Legend is same as (e). (g) Venn diagram overlap between proteins correlated with Oligo in our dataset, M2 Oligo proteome module from (e) and Oli0 marker transcripts from (f). Numbers indicate the total number of proteins in each overlapping set and selected protein names are shown. (h) Transcript log_2_ Fold Change values Oli0 population from Mathys et al. (2019) for markers of the Oligo cluster.

**Supplementary Figure 3.**
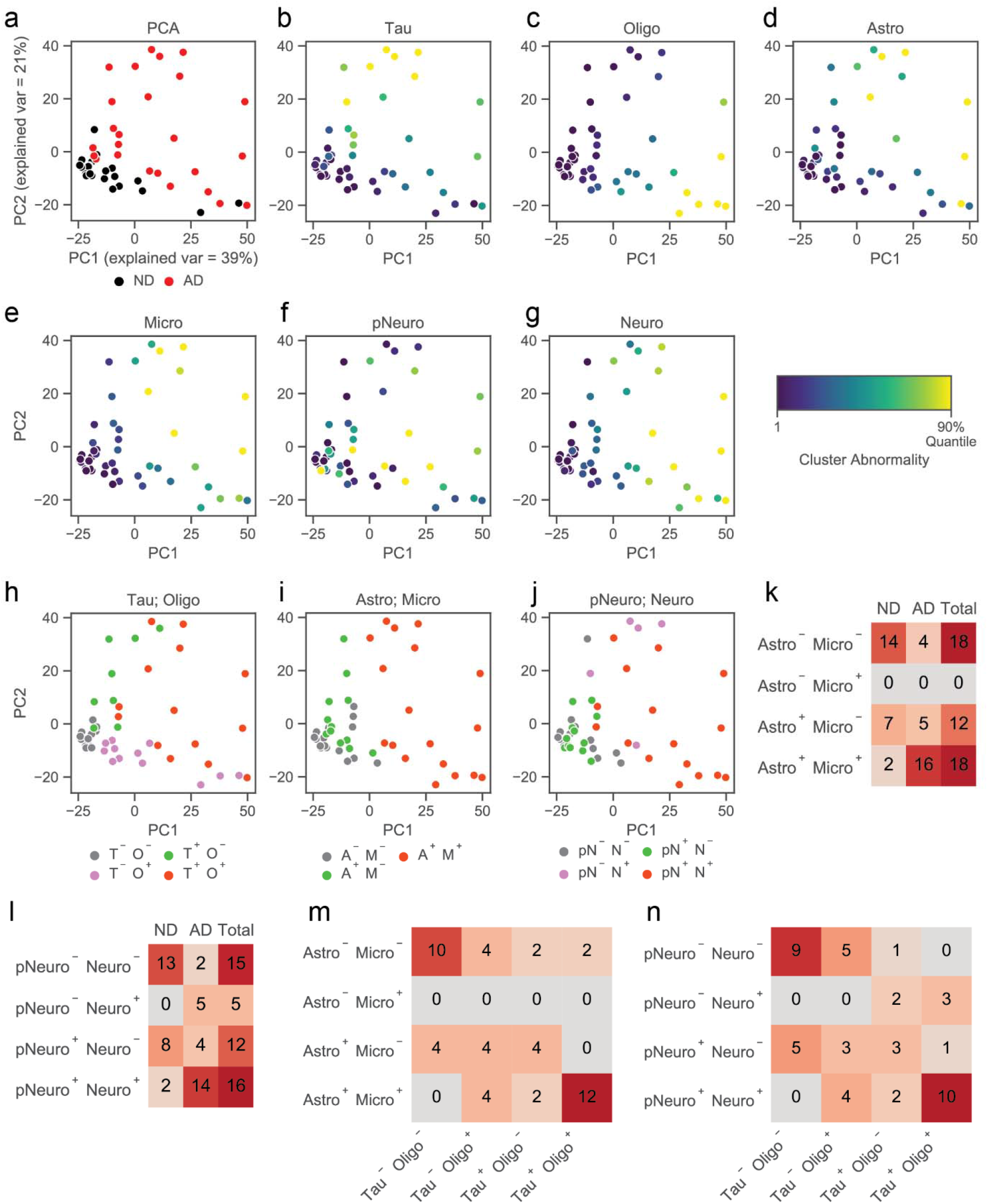
Peptide principle component analyses. (a) Principal component analysis (PCA) of peptides quantified in all samples. Patients are colored by (a) AD or ND status, (b) Tau, (c) Oligo, (d) Astro, (e) Micro, (f) pNeuro, and (g) Neuro cluster centroid values. (h-j) PCA of samples colored by patient stratifications using (h) Tau and Oligo, (i) Astro and Micro, and (j) pNeuro and Neuro cluster centroids. (k-l) Patients numbers for (k) Astro;Micro or (l) pNeuro;Neuro patient stratifications. (m-n) Overlap between Tau;Oligo and (m) Astro;Micro or (n) pNeuro;Neuro patient stratifications.

**Supplementary Figure 4.**
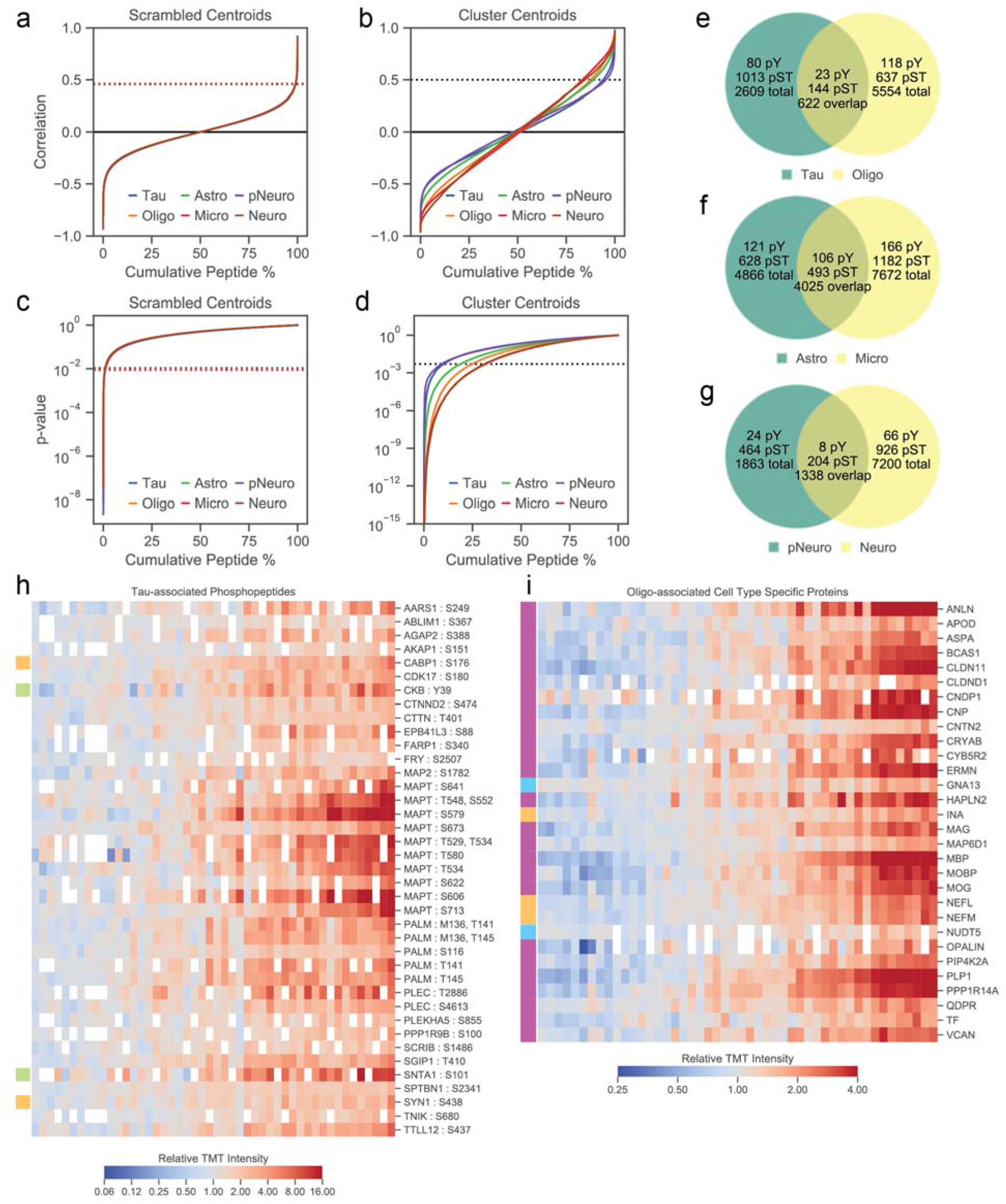
Peptide correlation analyses. (a-b) Distribution of Spearman’s correlation coefficient for peptides identified in ≥15 patients with (a) scrambled cluster centroids or (b) true cluster centroids. (c-b) Distribution of Spearman’s correlation p-values for peptides identified in ≥15 patients with (c) scrambled cluster centroids or (d) true cluster centroids. (e-g) Venn diagram showing the number of correlated peptides with (e) Tau and Oligo, (f) Astro and Micro, and (g) pNeuro and Neuro cluster centroids. Plots include the total number of associated peptides for each cluster, as well as the number of overlapping peptides between similar clusters. (h) Heatmap showing phosphopeptide fold changes for peptides that are significantly correlated with Tau cluster centroid. Samples are ordered by centroid values. Left color bars indicate proteins that were predicted to be cell-type specific. Green = Astrocyte, Purple = Oligodendrocyte, Yellow = Neuron, Blue = Microglia. Missing values are indicated as white boxes. (i) Heatmap showing fold changes cell-type specific proteins that are significantly correlated with Oligo cluster centroid. Samples are ordered by centroid value. Left color bar legend is same as (h).

**Supplementary Figure 4.**
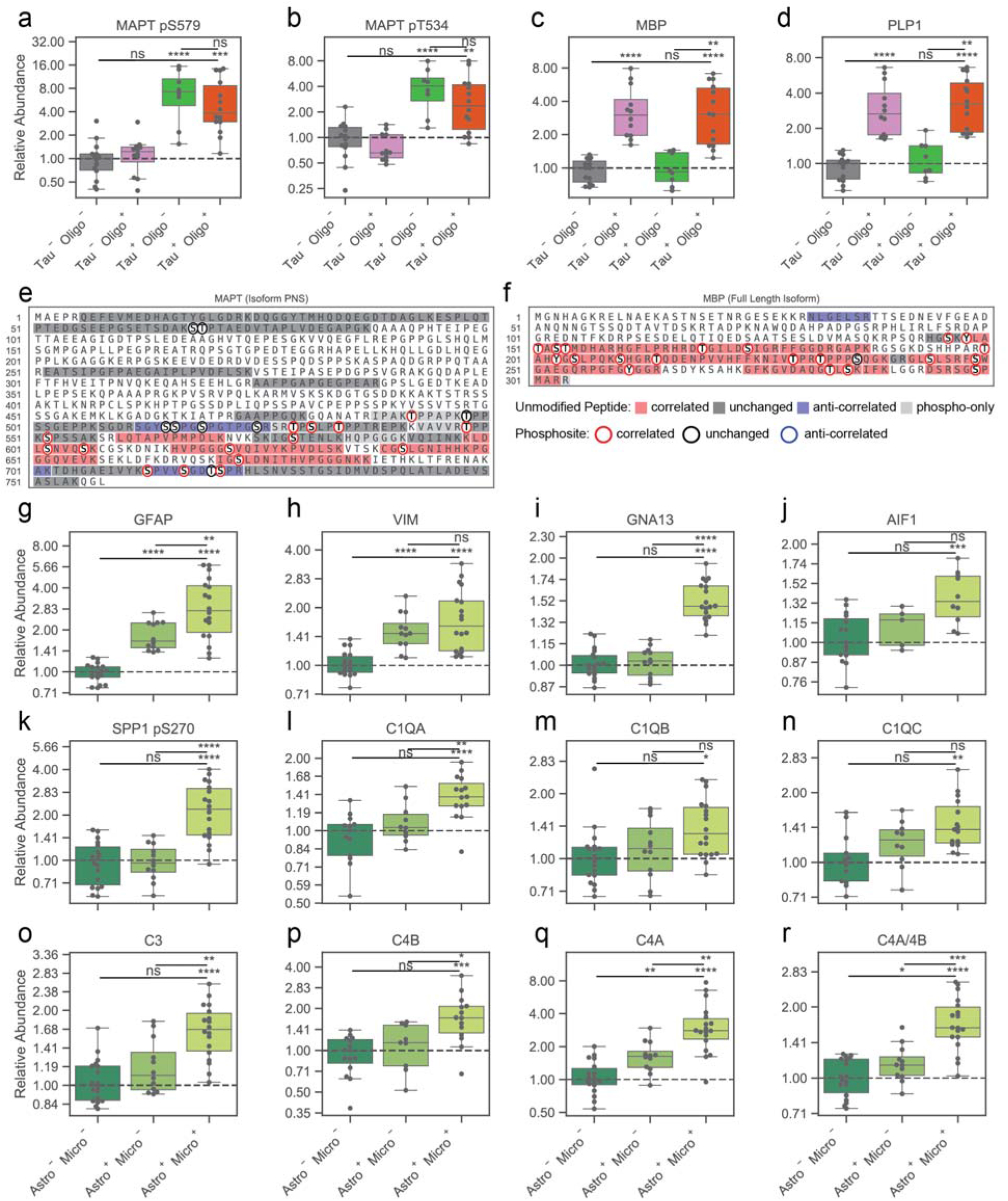
Analyses of peptide partial cluster correlation. (a-d) Fold changes for (a) MAPT pS579, (b) MAPT pT534, (c) MBP protein, and (d) PLP1 protein. Protein levels were estimated from the median abundance of all unmodified peptides that map uniquely to that protein. Values are normalized to the median of Tau^−^ Oligo^−^ patients. ns, not significant (p > 0.05); * p < 5e-2, **p < 1e-2; ***p < 1e-3; ****p < 1e-4. (e) All peptides mapping to the peripheral nervous system (PNS) isoform of MAPT. Colored bars indicate directional correlation for non-phosphorylated peptides. Red = correlated, blue = anti-correlated, grey = uncorrelated, light-grey = only phosphopeptides were seen in that region. Colored circles indicate phosphorylation sites that were quantified. Red circle = correlated, blue circle = anti-correlated, black circle = uncorrelated. (e) All peptides mapping to the full-length isoform of MBP. Legend is same as (e). (g-l) Fold changes for (g) GFAP protein, (h) VIM protein, (i) GNA13 protein, (j) AIF1 protein, (k) ITGB2 protein, and (l) SPP1 pS270 across Astro and Micro groups

**Supplementary Figure 6.**
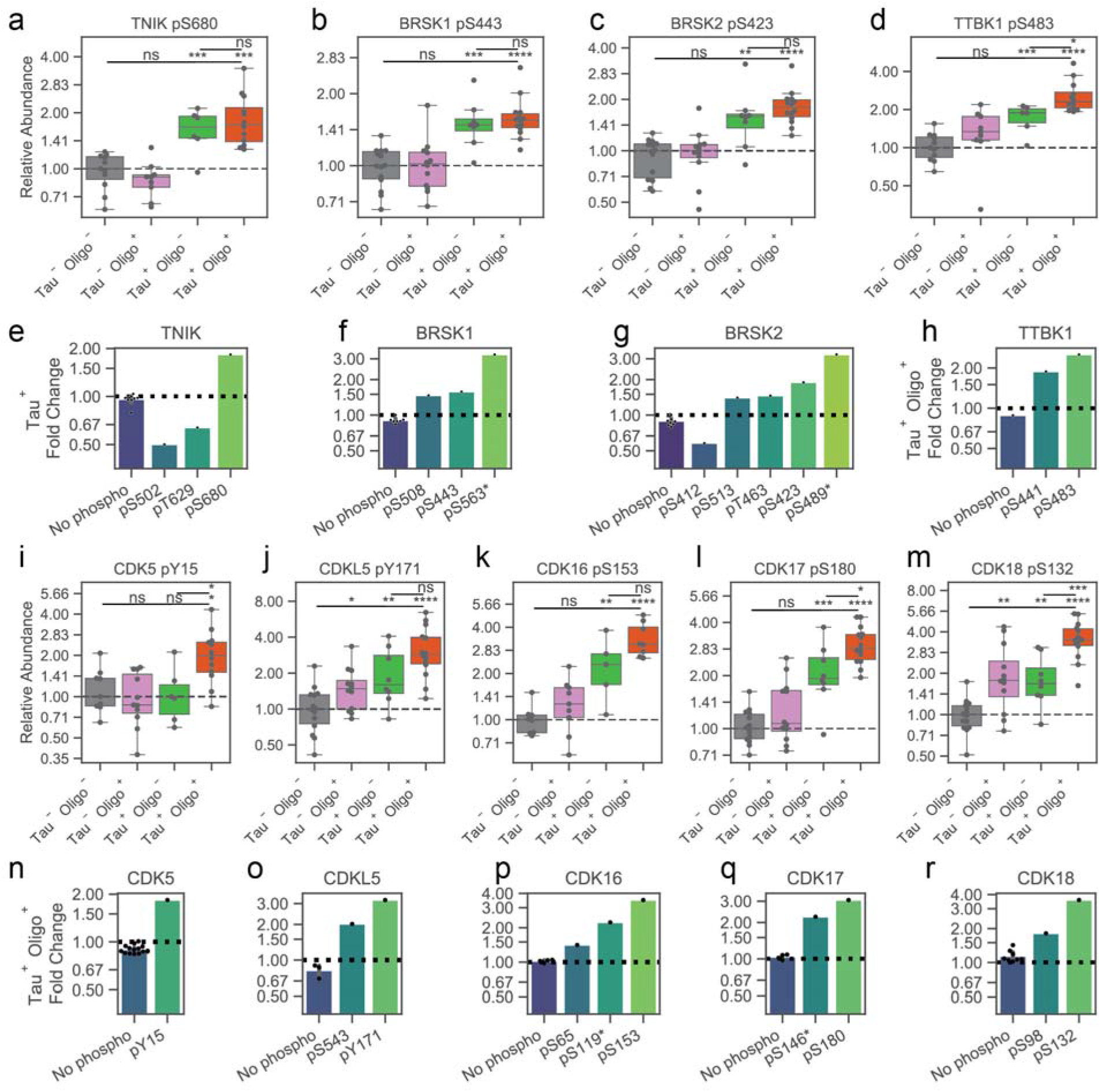
Tau kinase and CDK phosphosite abundances. (a-d) Fold changes for (a) TNIK pS680, (b) BRSK1 pS443, (c) BRSK2 pS423, and (d) TTBK1 pS483 across Tau and Oligo groups. (e-h) Fold changes for all non-phospho peptides and significantly changing phosphosites on (e) TNIK, (f) BRSK1, (g) BRSK2, (h) TTBK1. Each dot indicates one unique peptide. Phosphosites with ‘*’ are on peptides that map ambiguously to more than one protein. (i-m) Fold changes for (i) CDK5 pY15, (j) CDKL5 pY171, (k) CDK16 pS153, (l) CDK17 pS180, and (m) CDK18 pS132 across Tau and Oligo groups. (n-r) Fold changes for all non-phospho peptides and significantly changing phosphosites on (n) CDK5, (o) CDKL5, (p) CDK16, (q) CDK17, and (r) CDK18. Legend is same as (e-h).

**Supplementary Figure 7.**
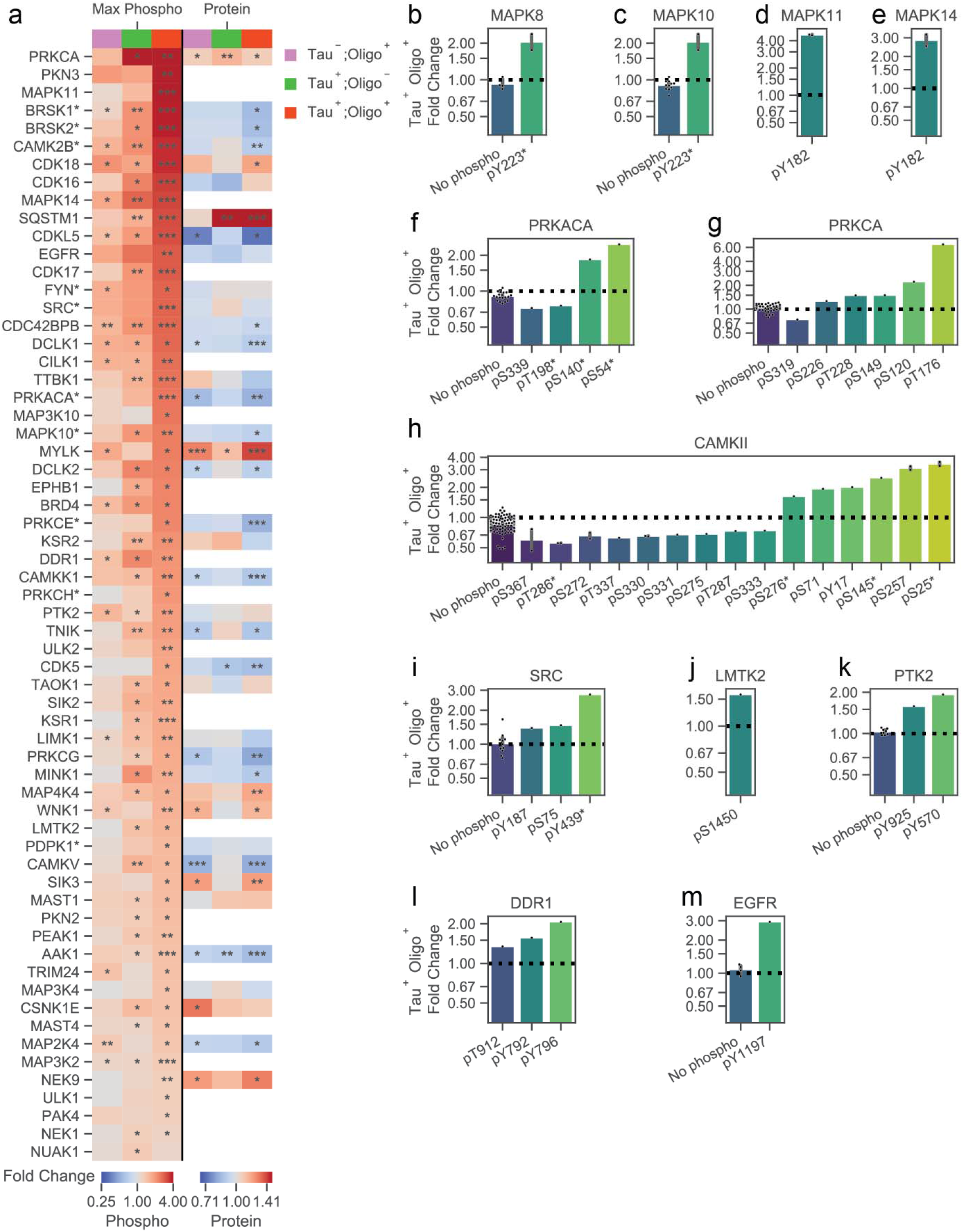
Accessory kinome analyses. (a) Maximum phosphosite fold changes and median protein abundances for all kinases shown in **Figure 4d**. Changes are calculated for each Tau;Oligo group compared to Tau^−^;Oligo^−^. *p < 5e-2; **p < 1e-3; ***p < 1e-4. (b-m) Fold changes for all non-phospho peptides and significantly changing phosphosites on (b) MAPK8 (c) MAPK10, (d) MAPK11, (e) MAPK14, (f) PRKACA, (g) PRKCA, (h) all CAMKII subunits, (i) SRC, (j) LMTK2, (k) PTK2, (l) DDR1, and (m) EGFR. Legend is same as **Supplementary Figure 6e**.

**Supplementary Figure 8.**
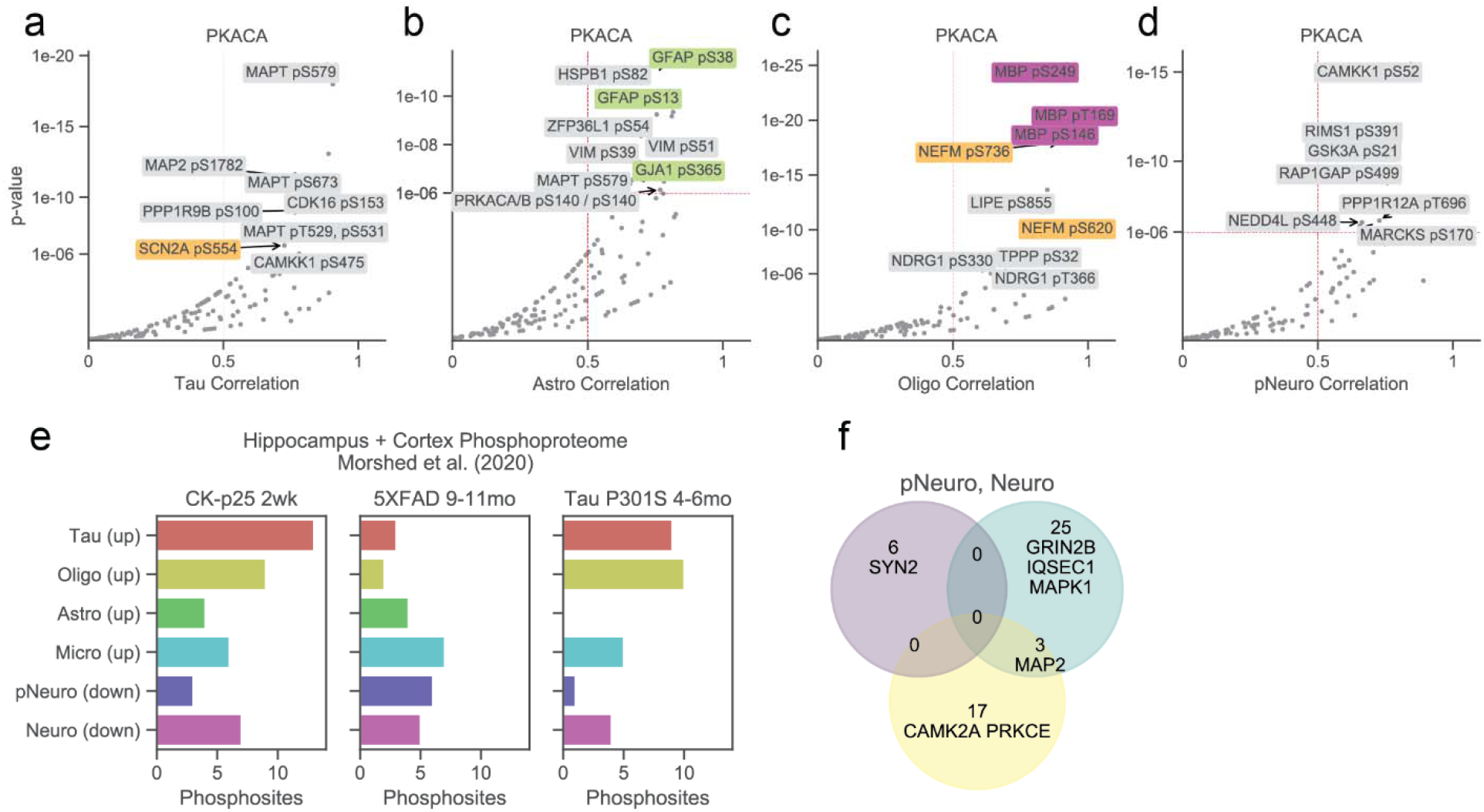
Kinase-substrate and AD mouse model analyses. (a-d) Volcano plots showing Spearman’s correlation coefficient and p-values between PKACA substrates and (a) Tau, (b) Astro, (d) Oligo, and (d) pNeuro cluster centroids. Labels are colored using the same scheme as **Supplementary Figure 4h**. (e) Number of phosphosites from AD mouse models identified in three mouse models of neurodegeneration that could be translated to human and were significantly correlated with each cluster centroid and shared directional changes. (f) Phosphoproteins that were identified to have at least one shared phosphosite with three mouse models of neurodegeneration. Proteins are shown as a Venn diagram between downregulated phosphosites and pNeuro or Neuro cluster centroids.

**Supplementary Figure 9.**
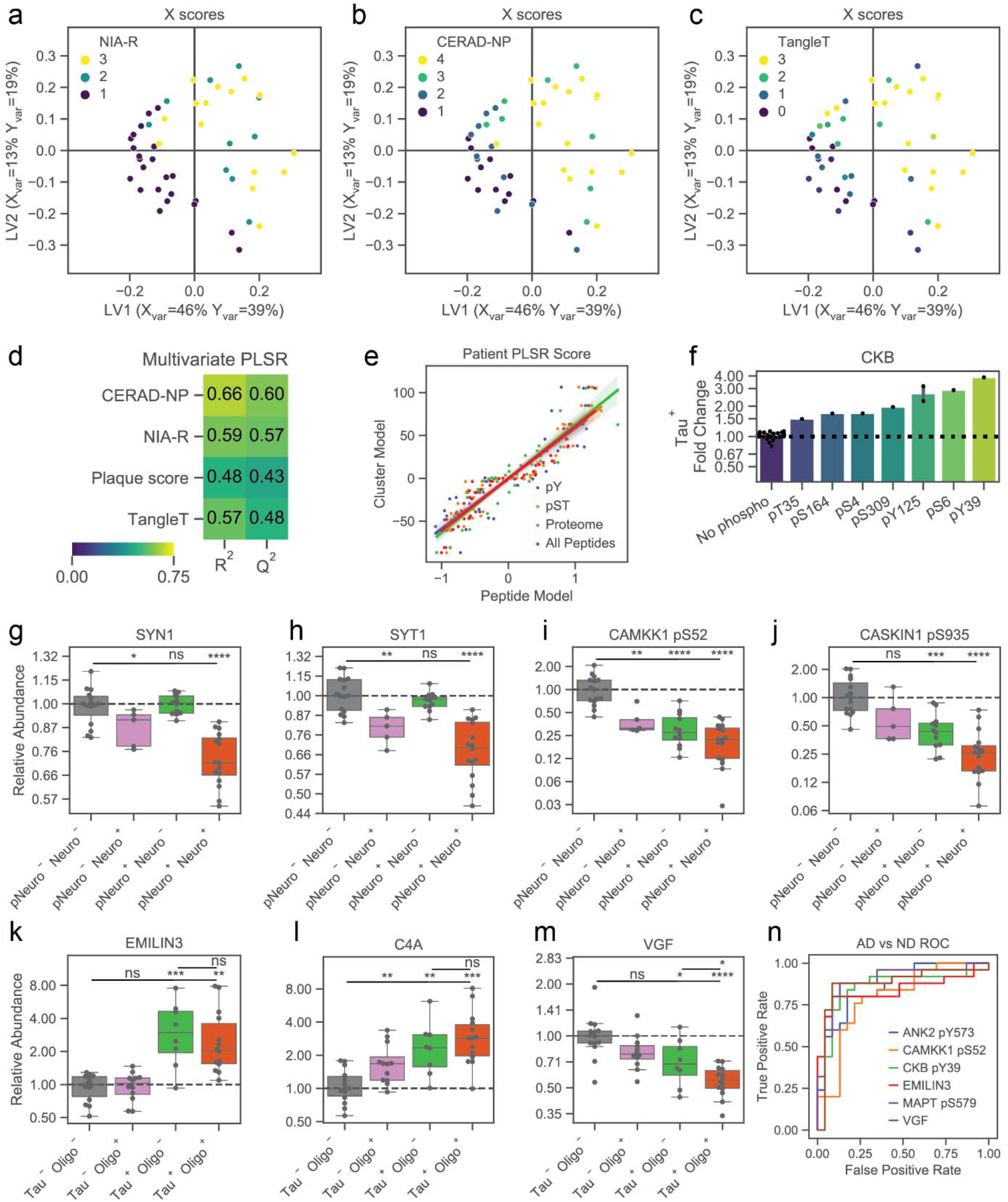
PLSR accessory analyses. (a-c) Latent variable plots for multivariate PLSR analysis for X scores vectors. Points are colored by patient (a) NIA-Reagan score, (B) CERAD-NP score, and (c) TangleT score. (d) K-fold validation of the multivariate PLSR model for all included histopathology variables. R^2^ and Q^2^ values are shown for training and test data predictions respectively. (e) Scatter plot showing PLSR patient scores generated from cluster centroids compared with subsets of the matrix of peptides quantified in all patients. (f) Fold changes for all non-phospho peptides and significantly changing phosphosites on CKB. Legend is same as **Supplementary Figure 6e**. (g-j) Fold changes for (g) SYN1, (h) SYT1, (i) CAMKK1 pS52, and (j) CASKIN pS935 across pNeuro and Neuro groups. (k-m) Fold changes for (k) EMILIN3, (l) C4A, and (m) VGF across Tau and Oligo groups. (n) ROC curve predicting AD status from phosphosites and protein highlighted in **Figure 5f-5h**.

